# Long-timescale dynamics shape visual conscious awareness

**DOI:** 10.64898/2026.09.03.748970

**Authors:** Zefan Zheng, Darinka Trübutschek, Yongchun Cai, Lucia Melloni

## Abstract

Consciousness is mostly studied as if it were composed of independent snapshots, aggregating reports over trials and experimental contexts, disregarding any temporal aspect. However, almost nothing is known about how conscious experience spontaneously fluctuates in time. To shed light on the temporal dynamics of conscious awareness, we evaluated the temporal autocorrelation of subjective visibility/confidence reports and objective performance across 20 datasets (N= 335). Two widely used measures of consciousness dissociate in temporal autocorrelation under full spatial attention: subjective measures followed an exponential decay function with long timescales (> 5s), whereas objective accuracy exhibited a slow linear decay function. But both measures share a common underlying temporal fluctuation dynamics, suggesting that the strong autocorrelation in subjective measures cannot be fully attributed to response bias or criterion shift, but might reflect a genuinely aperiodic fluctuation of conscious awareness. Causal transcranial magnetic stimulation (TMS) suggests that this temporal dependency of subjective measures is read out in the dorsolateral prefrontal cortex (DLPFC), where a similarly long neural timescale has been demonstrated. Together, these findings establish a temporal constraint for neural models of conscious access: candidate mechanisms should explain not only whether a stimulus becomes consciously accessible, but also how the probability of conscious access evolves over time.

## Introduction

Conscious perception is usually studied as if it were a sequence of statistically independent snapshots. In a typical experiment, a stimulus is presented, a participant reports whether it was seen, and responses are averaged across stimulus strengths, attentional states or experimental conditions^1–10^. This approach has been central to the search for the neural correlates of consciousness: by contrasting seen with unseen trials, researchers aim to isolate the mechanisms that determine whether sensory information becomes subjectively available^3,11–13^. Yet this snapshot view leaves open a basic question. When two physically similar stimuli are presented seconds or minutes apart, are the corresponding experiences independent events, or is the current state of awareness associated with its recent past?

There are several reasons to question the assumption of independence. Studies on perception, attention, memory and decision-making have documented systematic effects of previous stimuli and judgments on later reports^14–20^, an effect known as hysteresis, serial dependence or recalibration^21^. Most work, however, has focused on biases in perceptual content or decision history. Far less is known about whether subjective availability itself—the experience that a stimulus is visible, clear or confidently perceived—has its own temporal organization.

Yet the possibility that perceptual sensitivity may slowly fluctuate over time was already raised more than 70 years ago. Verplanck and colleagues showed, in a single observer, that successive responses to repeated near-threshold visual stimuli were not statistically independent^22^. Later work linked ongoing infraslow neural activity to slow fluctuations in detection performance^23–25^, and prestimulus neural states have been shown to predict whether near-threshold visual stimuli subsequently reach awareness^23,26,27^. Others found that participants can learn to increase perceptual sensitivity over time^8,28–31^. However, recently a series of studies related to confidence leak suggest that the serial dependence in subjective confidence responses is not perceptual but largely due to response bias and criterion shift^14,22,32–35^, as they found correlations in subjective confidence ratings across different stimuli and tasks within a trial or across adjacent trials. Therefore, what remains unclear is whether the serial effects in subjective reports are truly perceptual or merely post-perceptual, what the boundary condition of this serial dependence is, how long its influence persists, and whether common temporal dependencies are present in objective performance.

Investigation into objective performance is critical because visibility reports, confidence judgments, and objective performance do not necessarily reflect the same underlying processes. This dissociation is perhaps best illustrated by blindsight patients, and by blindsight-like phenomena in neurologically healthy observers, who can perform above chance while reporting little or no subjective awareness of the stimulus^36–40^. Comparing the temporal organization of subjective and objective measures therefore provides a means of testing whether they are governed by common or distinct slowly varying processes.

Further, recent work has revealed a systematic gradient of intrinsic neural timescales across the cortex^41,42^, with short timescales in sensory cortex (< 1s) and progressively longer timescales in association and prefrontal cortex (> 5s). These findings motivate the broader question of whether subjective and objective reports exhibit a reproducible temporal organization that neural models could ultimately explain, thereby providing additional constraints on ongoing debates regarding the relative contribution of prefrontal, associative, and sensory cortices to conscious perception^43,44^.

Here, building on these findings, we asked whether there is a lawful temporal structure governing the relationship between previous experience and current experience, and if so, whether that structure is shared or independent from those governing objective performance, and how long the timescale of such fluctuation is. To determine the systematicity of any temporal effect affecting subjective visibility and/or objective performance we re-analyzed 20 existing datasets spanning different paradigms, stimulus classes, visibility manipulations, response formats and subjective measures. Across these multi-lab datasets, participants provided visibility or confidence reports together with objective discrimination responses, allowing us to compare subjective and objective autocorrelation functions within the same behavioural sequences. Across datasets, subjective reports exhibited robust long-range temporal dependence, whereas objective performance showed substantially weaker temporal structure. These findings indicate that perceptual reports are not independent snapshots and provide a framework for investigating the slowly evolving internal states against which perceptual access unfolds. These temporal dynamics provide an additional behavioral constraint that mechanistic accounts of conscious access should explain. Beyond accounting for why identical stimuli are consciously perceived on some trials but not others, successful neural models should also explain how the probability of conscious access evolves systematically over time.

## Results

### Subjective and objective perceptual reports exhibit distinct temporal dependencies

We first asked whether subjective visibility and objective performance exhibit temporal dependencies and, if so, whether these dependencies are governed by similar or distinct temporal dynamics. We re-analyzed an existing dataset^8^ where participants judged both the visibility and identity of masked words presented under continuous flash suppression. Stimuli were shown at three contrast levels determining three visibility levels (high, medium, low; visible on 92%, 47% and 14% of trials, respectively), arranged into three block types: baseline blocks mostly containing liminal trials, visible-context blocks dominated by supraliminal trials, and invisible-context blocks dominated by subliminal trials (Fig. 1a,b). Because temporal dependencies can only be estimated when behavioral responses vary, all analyses were restricted to liminal trials, thereby avoiding the ceiling and floor effects present in supraliminal and subliminal conditions. For each participant, block type and trial lag (defined as the number of intervening liminal trials), we computed the autocorrelation separately for subjective visibility reports and objective performance by correlating all pairs of responses separated by the same lag (Fig. 1c). Baseline blocks were analyzed separately from visible- and invisible-context blocks because they contained substantially more liminal trials (94 vs. 20), allowing more reliable estimation of long-range temporal dependencies.

**Fig. 1.**
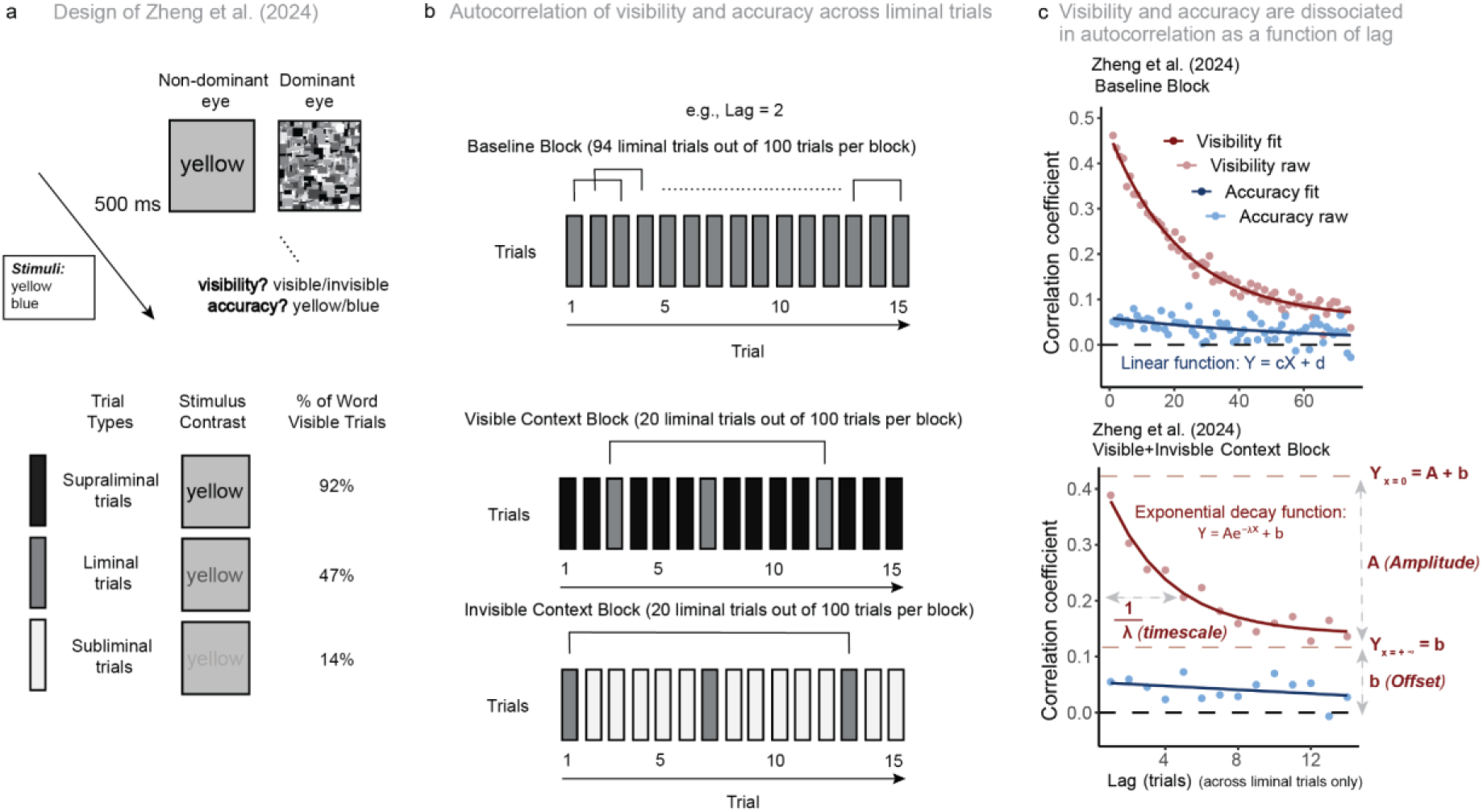
Autocorrelation functions for visibility and accuracy in Zheng et al.^8^. **a**, Experimental paradigm (adapted from Zheng et al.^8^). Participants viewed near-threshold words presented under continuous flash suppression and reported both subjective visibility (visible/invisible) and objective performance (word identity yellow/blue) on each trial. The experiment included supraliminal, liminal, and subliminal trials achieved by varying the word’s physical contrast, represented by filled colors. **b**, Illustration of the autocorrelation analysis. Autocorrelation coefficients were computed separately for subjective visibility reports and objective performance as a function of trial lag using liminal trials only. Analyses were performed for the baseline condition and for blocks in which liminal trials were embedded within predominantly visible or predominantly invisible contexts. **c**, Representative results from the Zheng et al. dataset. In the baseline condition (top), visibility reports were best modeled by an exponential decay function (in red), while objective performance coefficients follow a linear model (in blue) across successive trial lags. In the visible and invisible context blocks (bottom), the autocorrelation of visibility reports was well described by an exponential decay, whereas objective performance showed little evidence of systematic temporal dependence. In the exponential decay function, the parameter “b” represents the offset, and “A” the amplitude. “A + b” indicates the maximal coefficient when the lag approaches 0. The parameter “λ” denotes the decay rate constant, with its inverse corresponding to the lag at which the coefficients drop to 1/e (approximately 0.367) times of “A”.

We found that both subjective visibility and objective performance exhibited clear temporal dependency (Fig. 1c), demonstrating that neither measure behaved as a sequence of statistically independent observations. Importantly, the temporal profile differed between measures: Subjective visibility was consistently better described by an exponential decay than by a linear model, both in baseline blocks (F(1) = 78.281, P < 0.001, w_AICc_linear_ = 0, w_AICc_exponential_ = 1) and in visible/invisible-context blocks (F(1) = 10.582, P = 0.001, w_AICc_linear_ = 0.014, w_AICc_exponential_ = 0.986). By contrast, objective performance was adequately captured by a linear function (baseline: F(1) = 0.306, P = 0.580, w_AICc_linear_ = 0.703, w_AICc_exponential_ = 0.297; visible/invisible context: F(1) = 0.002, P > 0.999, w_AICc_linear_ = 0.733, w_AICc_exponential_ = 0.267), indicating a shallower, approximately linear history dependence. Thus, while both subjective and objective reports exhibited temporal dependence, only subjective visibility displayed a characteristic temporal timescale.

The exponential model of subjective visibility was parameterized by an initial amplitude (A), an asymptotic offset (b), and a decay constant (λ), whose inverse defines the characteristic timescale of the temporal dependency. We further estimated the temporal response window (TRW) as the lag at which the autocorrelation approached its asymptote (defined as 2% of the initial amplitude). Given that each trial lasted at least 3 s, both the estimated characteristic timescale and the temporal response window exceeded 50 seconds, indicating that the influence of previous perceptual states extended across dozens of successive perceptual decisions. Interestingly, this behavioral timescale substantially exceeded intrinsic timescales typically reported for posterior areas using electrophysiological recordings and computational modelling^42,45–48^. Specifically, Chaudhuri^41^ observed temporal autocorrelation of neuronal firing with timescales over 5 seconds only in the macaque prefrontal cortex. Although behavioral and neural timescales are not directly comparable, this correspondence suggests that the observed temporal dynamics are unlikely to arise solely from rapidly fluctuating sensory processes.

Objective performance, by contrast, was well captured by a linear autocorrelation on lag. In baseline blocks, the slope was small but significantly negative (*β* = −0.0005, SE < 0.001, P < 0.001) and the intercept significantly positive (*β* = 0.055, SE = 0.006, P < 0.001), indicating a slow, monotonic decline from a positive starting value. In visible/invisible-context blocks, the slope estimate was similar but no longer significant (*β* = −0.0003, SE < 0.001, P = 0.274), likely reflecting reduced power, whereas the positive intercept remained robust (*β* = 0.055, SE = 0.013, P < 0.001). The presence of a long, approximately linear decay in objective performance is inconsistent with a purely criterion-shift explanation of the exponential temporal dependence observed for visibility. Instead, both measures appear to be influenced by slowly evolving internal states, although exhibiting qualitatively different temporal dynamics in subjective and objective reports.

### Strong generalizability for fully attended foveal stimuli, with attenuation in divided-attention and peripheral paradigms

As the dissociation observed in Fig 1 between an exponential autocorrelation in visibility and a shallow, nearly linear, dependence in objective performance could represent an idiosyncrasy of the original paradigm, we next evaluated its generalization across tasks, stimuli and laboratories, by re-analyzing sixteen datasets: thirteen from the Confidence Database^56^, one from Sand & Nilsson^51^ and two from Zheng et al.^8^. To stress test the finding we evaluated different visibility manipulations (backward masking, iconic memory, low contrast), stimuli (letters, words, faces, objects colored disks, Gabor patches), subjective measures (visibility and confidence), objective readouts (2 alternative forced choice and continuous estimation), foveal and peripheral presentations, and trial durations between ∼3–10 s. Rating formats ranged from binary to four-level scales for visibility and two- or three-level scales for confidence. We applied the same autocorrelation and model-comparison pipeline as in the first dataset (Fig. 2).

**Fig. 2.**
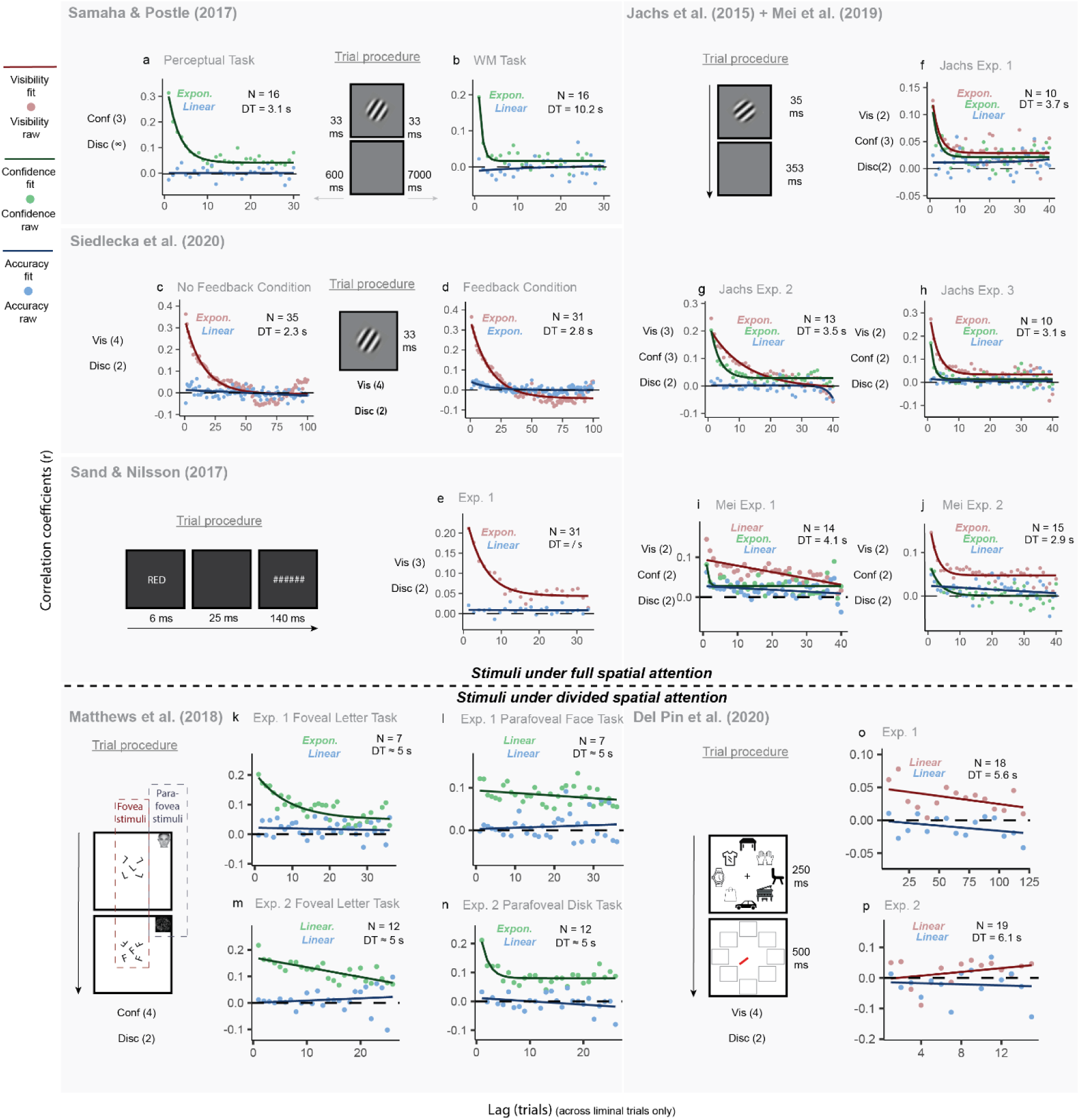
Distinct temporal dependencies of subjective and objective perceptual reports generalize across experimental paradigms. To determine whether the dissociation observed in Fig. 1 generalizes beyond a single experimental paradigm, we reanalyzed previously published datasets spanning a broad range of perceptual tasks, subjective report measures, stimulus classes, and experimental manipulations. Across all studies, participants performed at least one subjective perceptual judgment (visibility and/or confidence) together with an objective discrimination task on briefly presented stimuli. The upper panels illustrate the paradigms using low-contrast Gabor patches under working-memory delays (Samaha & Postle^49^ in a and b), masked Gabor stimuli with or without discrimination feedback (Siedlecka et al.^50^ in c and d), backward-masked colour words (Sand & Nilsson^51^ in e), and backward-masked Gabor stimuli with different subjective and objective response formats (Jachs et al.^52^; Mei et al.^53^ in f, g, h, i, and j). The lower panels show paradigms requiring divided spatial attention, including whole-report tasks with concurrent peripheral discrimination (Matthews et al.^54^ in k, l, m, and n) and iconic-memory paradigms using multi-object displays (Del Pin et al.^55^ in o and p). The dashed horizontal line separates experiments in which subjective and objective judgments referred to the same foveal stimulus from those requiring divided spatial attention between central and peripheral stimuli. For each dataset, autocorrelation functions were computed separately for subjective visibility (red), confidence (green), and objective performance (blue). Dots represent empirical autocorrelation coefficients across trial lags, and curves indicate the best-fitting model. *Expon.* (i.e., exponential decay function) or *Linear* (i.e., linear function) denotes the best-fitting mathematical function to the curve of the corresponding color. For visual comparison, exponential curves are displayed for all datasets, even when the linear model was statistically preferred. Sample size (N) and mean trial duration (DT) are shown in the upper right corner of each panel, when available. Across highly diverse paradigms, subjective reports (visibility and confidence) consistently exhibited stronger, typically exponential, temporal dependencies than objective performance, demonstrating that the dissociation identified in Fig. 1 generalizes across perceptual tasks, response modalities, and experimental contexts.

Despite these substantial differences in experimental design, the temporal organization of subjective reports proved remarkably consistent. We found that subjective reports showed a highly consistent exponential profile for foveally and fully attended stimuli: Visibility was better fit by an exponential than a linear model in seven of eight datasets (Fig. 2c–h,j; Supplementary Table 1). Confidence showed the same pattern across all seven datasets in which it was measured (Fig. 2a,b,f–j; all F(1) > 5.232, Ps < 0.023). By contrast, exponential decay was rarely observed when spatial attention was divided or stimuli appeared in the periphery: only two of six such datasets showed significant exponential components in the subjective measures (Fig. 2k,n).

Objective performance displayed a markedly different profile. In thirteen of sixteen datasets, the exponential model failed to outperform the linear model despite the former having one additional parameters (Fig. 2a–c,e–j,k–m; all F(1) < 3.837, Ps > 0.05; Supplementary Table 1). Three apparent exceptions had specific explanations: Two (Fig. 2n, p) exhibited exponential fits driven primarily by a small number of observations at the longest lags, and one dataset (Fig. 2d) was providing trial-by-trial feedback, suggesting that explicit feedback on performance can modify the function from linear towards an exponential. Overall, the dominant pattern across paradigms was an exponential profile for subjective measures and an approximately linear profile for objective accuracy, with the exponential pattern attenuated when attention was divided.

To quantify the stability of these temporal profiles, we performed cross-dataset random-effects synthesis of the fitted model parameters: amplitude *A*, decay constant *λ* and offset *b* for the exponential model, and slope *c* and intercept *d* for the linear model (Fig. 3). Across ten visibility datasets, exponential amplitudes were significantly greater than zero in eight datasets (Supplementary Table 2), positive decay constants in seven, and asymptotic offsets in six (A: *β* = 0.244, 95% CI 0.164–0.320, t₁₀ = 6.70, P < 0.001; T² = 0.013, I² = 94.6; λ: *β* = 0.098, 95% CI 0.038–0.158, t₁₀ = 3.59, P = 0.004; T² = 0.004, I² = 74.7%; b: *β* = 0.028, 95% CI −0.000–0.057, t₁₀ = 2.20, P = 0.050; T² = 0.002, I² = 96.3%, respectively). These cross-dataset meta-analytic estimates suggest that the behavioral timescale of subjective visibility (i.e., the inverse of decay constant λ) is 10.2 trials (a timescale estimate in clock time is not available, as some datasets did not record clock time; but see Fig. 7 for a timescale approximation in clock time).

**Fig. 3.**
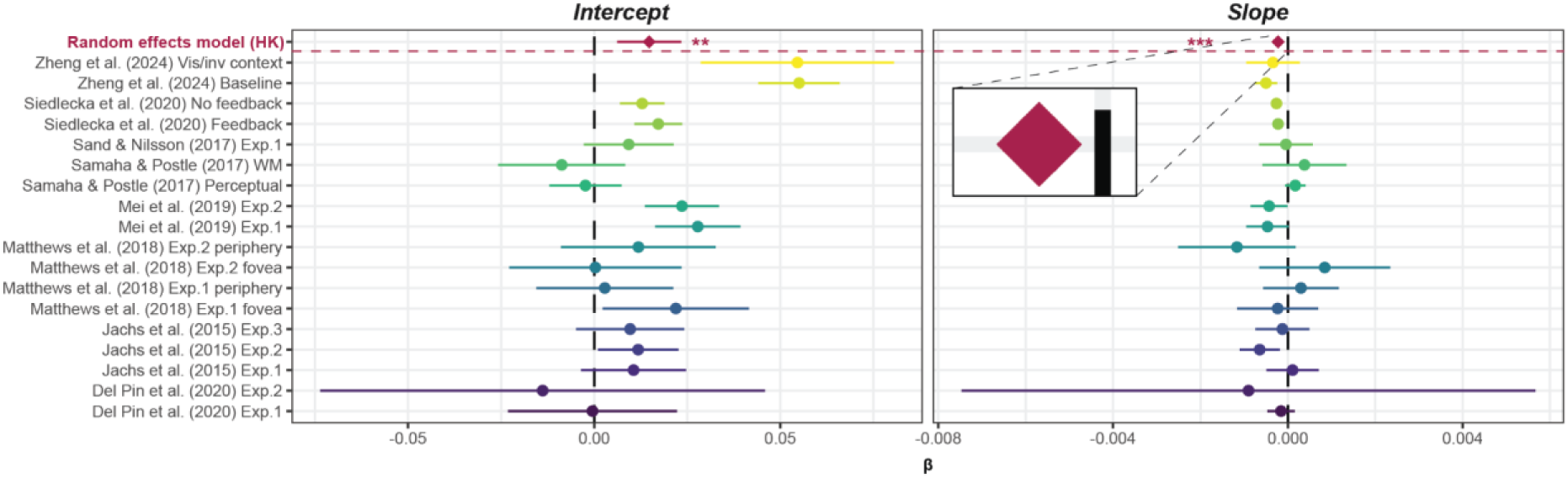
Cross-dataset random-effects synthesis of the autocorrelation parameters of objective performance. Forest plots show study-level estimates and 95% confidence intervals for the intercept (left) and slope (right) of the fitted autocorrelation function for objective performance. Each colored circle represents the identity of a given dataset; the maroon diamonds denote the pooled estimates from the Hartung–Knapp (HK) random-effects model. Vertical dashed black lines indicate zero effect. The pooled intercept was significantly positive, indicating a reliable overall positive autocorrelation in objective performance, whereas the pooled slope was significantly negative, indicating that this autocorrelation decayed with lag. Asterisks mark the significance of the pooled effects. Together, these results show that objective performance, although much weaker than subjective measures, nevertheless exhibits a small but reliable lag-dependent temporal dependence across datasets. * P < 0.05; ** P < 0.01; *** P < 0.001.

Confidence judgments exhibited an almost identical pattern, with significantly positive amplitudes, decay constants and offsets across studies (Supplementary Table 3), corresponding to a shorter characteristic timescale of approximately 3.0 trials (A: *β* = 0.193, 95% CI 0.118–0.268, t₁₀ = 5.74, P < 0.001; T² = 0.007, I² = 69.1%; λ: *β* = 0.328, 95% CI 0.160–0.496, t₁₀ = 4.34, P = 0.002; T² = 0.023, I² = 58.5%; b: *β* = 0.035, 95% CI 0.017–0.052, t₁₀ = 4.45, P = 0.001; T² = 0.001, I² = 94.6%, respectively).

Notably, amplitude and decay constants were not reliably different from zero in four datasets with divided spatial attention (confidence: Fig. 2l, m; visibility: Fig. 2o, p); only Experiment 2 of Matthews et al.^54^ showed a marginally positive amplitude (*β* = 0.237, t₂₃₁ = 2.06, P = 0.041).

Accuracy results were more heterogeneous. Eight of eighteen datasets exhibited significantly positive intercepts and five showed significantly negative slopes (Supplementary Table 4). Nonetheless, the cross-dataset random-effects synthesis of accuracy estimates revealed a reliable positive intercept (d: *β* = 0.015, 95% CI 0.006–0.023, t₁₆ = 3.62, P = 0.002; T² < 0.001, I² = 82.4%) and a small but significant negative slope (c: *β* = −0.0002, 95% CI −0.0003 to −0.0001, t₁₆ = −4.28, P < 0.001; T² < 0.001, I² = 41.5%; see Fig. 3), consistent with a weak linear decline in autocorrelation over lag (See Supplementary Figure S1 for cross-dataset random-effects synthesis of subjective measures).

Together, these results demonstrate that the dissociation identified in Fig. 1 is not specific to a particular task, stimulus class, laboratory, or subjective report format. Instead, visibility and confidence judgments generally exhibit robust exponential temporal dynamics when stimuli are presented foveally under focused attention, whereas objective performance follows a slower, approximately linear decay. The robust evidence for temporal dependence in objective performance across paradigms is hard to reconcile with a purely criterion-based explanation. Instead, the results are consistent with subjective and objective behavior being influenced, at least in part, by a slowly evolving latent state. A slowly varying perceptual threshold is one candidate implementation of such a state, but the present analyses do not uniquely identify its underlying nature. What remains open, however, is what computational mechanism could drive those temporal profiles.

### Aperiodic latent dynamics reproduce the temporal profiles of subjective and objective reports

At least two classes of mechanisms could, in principle, generate the observed temporal dependencies. Perceptual thresholds may fluctuate periodically, reflecting oscillatory neural dynamics, or they may evolve as a slowly varying aperiodic process driven by infraslow spontaneous activity^25,57^. These alternatives make qualitatively different predictions for the temporal organization of perceptual reports (Fig. 4). To test whether a perceptual threshold fluctuates periodically or aperiodically, we simulated threshold fluctuations using periodic and aperiodic functions, respectively, and computed their autocorrelation functions.

**Fig. 4.**
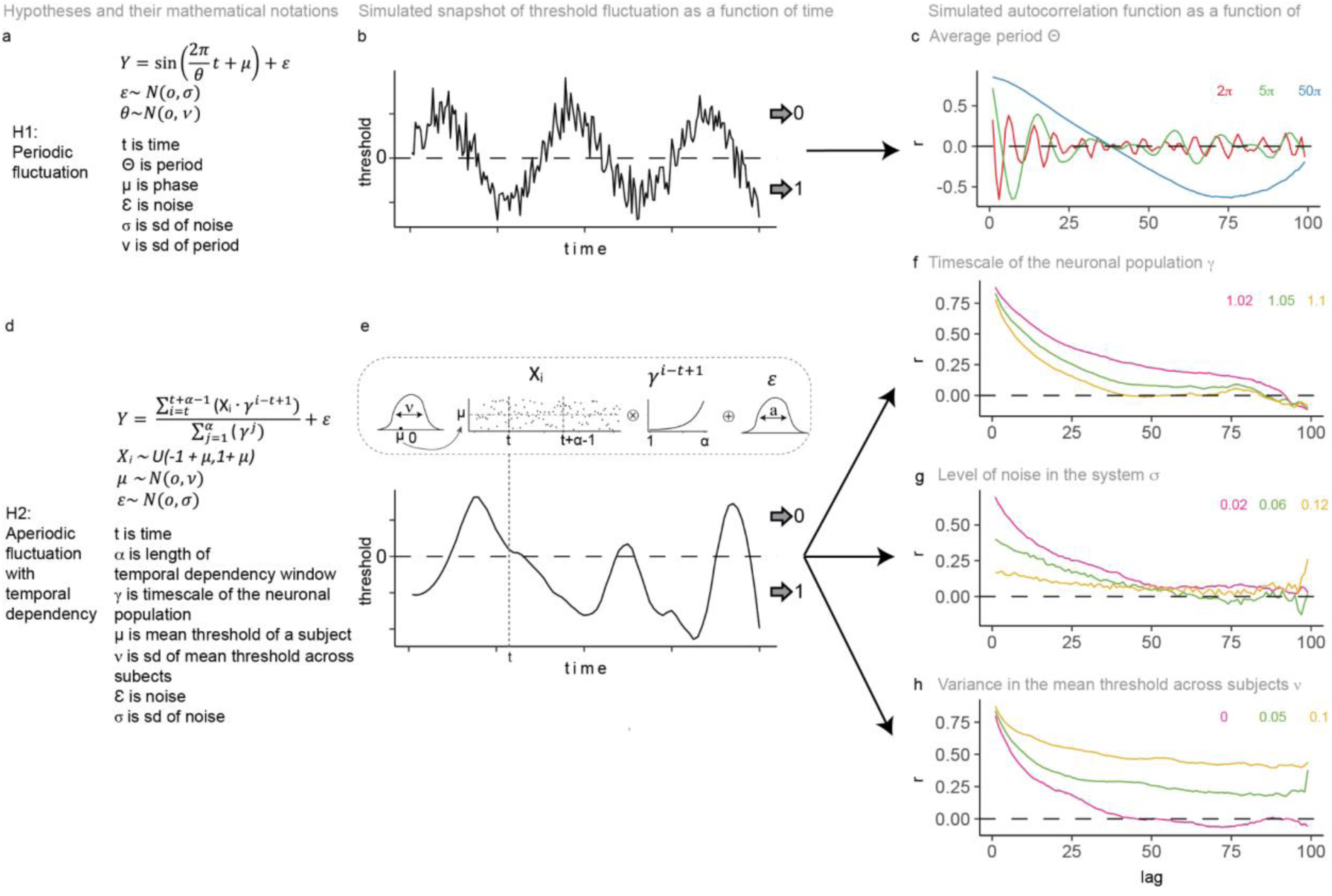
Simulating two candidate mechanisms for fluctuations in perceptual threshold. We simulated a periodic process (a-c), and an aperiodic process with exponential temporal dependency (d-h). For each hypothesis, panels show the model formulation (a, d), example threshold trajectories (b, e), and the resulting autocorrelation functions (c,f-h). Under Hypothesis 1, perceptual thresholds fluctuate periodically (a,b), producing oscillatory autocorrelation functions whose amplitude gradually decreases because of noise (c). Under Hypothesis 2, perceptual thresholds evolve as a slowly varying stochastic process generated by applying an exponentially weighted moving average (memory parameter γ) to white-noise input (d,e). The memory parameter γ determines the characteristic timescale of the behavioral autocorrelation (f), observation noise (σ) determines its amplitude (A; g), and between-subject variability in the mean threshold (ν) determines the asymptotic offset (b; h).

Under the periodic hypothesis (Hypothesis 1), perceptual thresholds oscillate around a stable mean with a characteristic period that varies across observers (Fig. 4a,b). This model predicts oscillatory autocorrelation functions, with alternating positive and negative correlations whose period increases with the underlying oscillation (Fig. 4c). Although such behavior can account for the few datasets in which autocorrelation briefly crosses zero (for example, Fig. 2c,d), it fails to reproduce the predominantly monotonic decay observed across experiments.

By contrast, the aperiodic hypothesis (Hypothesis 2) assumes that perceptual thresholds evolve as a long-memory stochastic process (Fig. 4d,e). We implemented this using an exponentially weighted rolling average of white-noise input, where the decay constant (γ) determines the persistence of the latent process. Unlike the periodic model, this simple mechanism naturally generates monotonic exponential autocorrelation functions resembling those observed empirically.

The model further revealed a straightforward relationship between latent dynamics and the observed autocorrelation profiles. The decay constant γ determined the characteristic timescale of behavior (Fig. 4f): slower latent dynamics yielded longer-lasting temporal dependencies. Increasing observation noise (σ) selectively reduced autocorrelation amplitude while preserving the underlying timescale (Fig. 4g). At sufficiently high noise levels, the resulting autocorrelation was empirically well approximated by the shallow, nearly linear profile observed for objective performance. Finally, variability in the mean threshold across observers (ν) determined the asymptotic offset of the autocorrelation function (Fig. 4h), with greater between-subject variability producing larger positive offsets.

Together, these simulations show that slowly varying aperiodic threshold fluctuations are sufficient to generate (i) exponential decay with large offsets, as observed for visibility and confidence (low γ, low σ, high ν), and (ii) shallow, quasi-linear profiles (low γ, high σ). The periodic model cannot account for this pattern. Our data are thus most consistent with inherently aperiodic fluctuations in perceptual thresholds whose decay constant reflects the timescale of underlying neural dynamics, with amplitude constrained by noise, and where the offset is determined by how the stimulus strength was stringently tailored to each observers’ liminal threshold. Thus, the distinct temporal profiles of subjective and objective reports need not arise from different latent processes but can emerge from the same underlying dynamics when measured with different levels of noise. These results are consistent with the proposal that infraslow aperiodic activity modulates conscious perception^57^.

### Temporal dynamics are not readily explained by simple response repetition and vary with elapsed time and measurement sensitivity

In the cross-dataset analyses above, all five fitted parameters (A, λ, b, c, d) were reliably different from zero, but also showed medium-to-high heterogeneity across datasets. To identify the sources of this heterogeneity and to probe the nature of the temporal profiles, we carried out a set of targeted, pairwise comparisons between datasets that differed in only one experimental factor (Fig. 5). Guided by our simulations, we interpreted changes in amplitude (A) as changes in measurement sensitivity and noise and changes in decay constant (λ) as differences in the timescale of the underlying neural populations. Offsets were set to zero for all datasets as between-study variance in offsets has a trivial explanation (different variances across individual thresholds). To equate designs with different densities of liminal trials, we expressed lag in standardized units reflecting average elapsed trials within a block (see **Methods**).

**Fig. 5.**
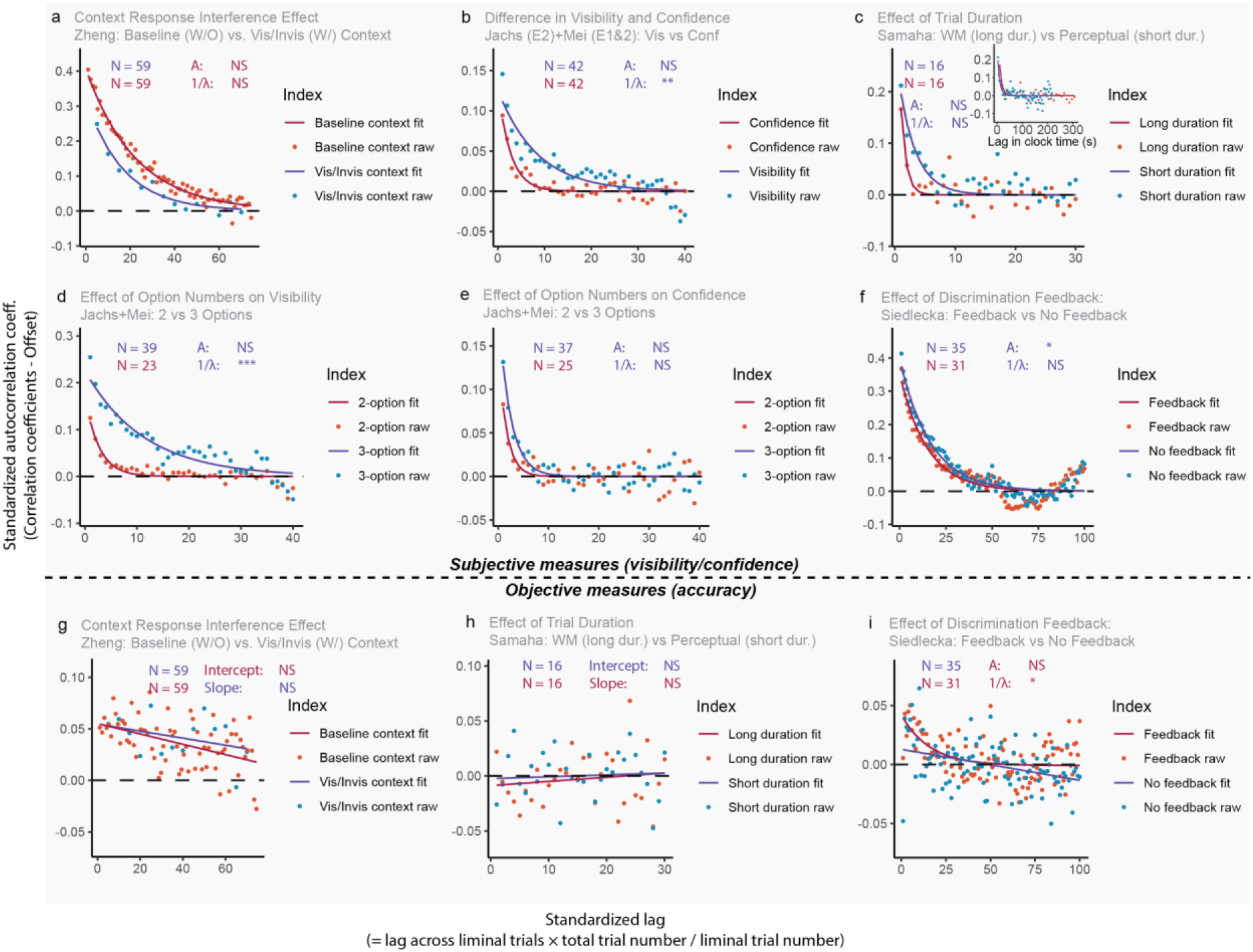
Measurement sensitivity and time rather than motor response shape temporal dependence on subjective and objective measures. Pairwise comparisons were performed between experimental conditions differing in a single factor. (a,g) Baseline versus visible/invisible context blocks (Zheng et al.^8^) testing response repetition. (b) Visibility versus confidence (Jachs et al.^52^; Mei et al.^53^) (c,h) Perceptual versus working-memory conditions (Samaha and Postle^49^), testing the effect of elapsed time. (d,e) Two versus three response levels (Jachs et al.^52^; Mei et al.^53^), testing measurement sensitivity. (f, i) No-feedback versus feedback conditions (Siedlecka et al.^50^) testing the effect of performance feedback. Sample sizes are indicated for each condition. Statistical comparisons are shown for the exponential model parameters (subjective measures) and the linear model parameters (objective performance), except in j, where the feedback condition was better fit by an exponential model. Parameter labels are coloured according to the condition with the larger estimated value. The x-axis shows standardized lag and the y-axis the autocorrelation coefficient (r). NS, not significant; P < 0.05; ** P < 0.01; *** P < 0.001.

A first possibility is that long-range temporal dependence in subjective measures reflects a tendency to repeat the same response. Zheng et al.^8^, provide a direct test of this hypothesis because in the visible and invisible context blocks, successive liminal trials were separated on average by four supraliminal or subliminal trials (Fig. 1b). If response bias were dominant, this interruption should strongly reduce the autocorrelation measured across liminal trials. It did not. In fact, autocorrelation parameters in context blocks did not differ significantly from those in baseline blocks (A: *β̂* = −0.060, SE = 0.070, P = 0.390; λ: *β̂* = 0.016, SE = 0.026, P = 0.528; Fig. 5a). This finding argues against a purely local sticky-response or motor-perseveration account.

If behavioral autocorrelations reflect intrinsic fluctuations in neural activity that modulate perceptual threshold, they should scale with elapsed time rather than with the number of trials. We tested this by comparing the perceptual and working-memory (WM) conditions in Samaha and Postle^49^, which differed only in trial duration (∼3 s versus ∼10 s). Expressed in trial units, the estimated timescales differed markedly (2.818 versus 0.883 trials; Fig. 5c). After converting to physical time, however, they were nearly identical (8.46 versus 8.83 s). Objective performance parameters did not differ between conditions (intercept: *β̂* = −0.008, SE = 0.012, P = 0.544; slope: *β̂* < 0.001, SE < 0.001, P = 0.617; Fig. 5h). These results support a time-based, rather than trial- or response-based, interpretation of the decay.

Our simulations further predicted that improving measurement sensitivity (i.e., signal-to-noise ratio in the readout system) should primarily increase the amplitude of the autocorrelation while leaving its characteristic timescale largely unchanged. We tested this prediction in two independent ways. First, one proposal is that increasing the number of alternatives in forced-choice tasks enhances the sensitivity of visibility reports^52,58,59^. Pooling five datasets from Jachs et al.^52^ and Mei et al.^53^, we compared 3AFC versus 2AFC versions of subjective measures. Visibility autocorrelations showed significantly larger amplitudes and somewhat slower decay in 3AFC than 2AFC (A: *β̂* = 0.100, SE = 0.035, P = 0.004; λ: *β̂* = −0.219, SE = 0.109, P = 0.044; Fig. 5d,e). Confidence showed a similar but non-significant trend (A: *β̂* = 0.053, SE = 0.049, P = 0.277; λ: *β̂* = −0.212, SE = 0.184, P = 0.249), demonstrating that more options in subjective measures do increase signal-to-noise ratio. Second, we examined the impact of trial-by-trial feedback, a common method for increasing the signal-to-noise ratio in objective discrimination. Using Siedlecka et al.^50^’s feedback and no-feedback blocks, which were otherwise identical, we found that feedback significantly increased the amplitude of the visibility autocorrelation (A: *β̂* = −0.046, SE = 0.022, P = 0.040) without changing its decay constant (λ: *β̂* = −0.003, SE = 0.007, P = 0.696; Fig. 5f). Linear fits to objective performance showed no feedback effect (intercept: *β̂* = −0.004, SE = 0.004, P = 0.333; slope: *β̂* = 3.65×10⁻⁵, SE = 7.70×10⁻⁵, P = 0.636). Remarkably, in the feedback condition, objective performance itself was better described by an exponential than a linear model, and its exponential decay constant was significantly larger than in the no-feedback condition (λ: *β̂* = 0.064, SE = 0.028, P = 0.024). Thus, increasing measurement sensitivity revealed temporal dynamics in objective performance that more closely resembled those of subjective reports, precisely as predicted by the model (Fig. 4i).

Finally, we asked whether different subjective reports are governed by the same temporal process. Pooling data from Jachs et al.^52^ and Mei et al.^53^ revealed that visibility decayed significantly more slowly than confidence (*β^* = −0.254, SE = 0.094, *P* = 0.007; Fig. 5b), indicating that the temporal integration underlying visibility extends over longer timescales than that supporting confidence judgments.

Across all contrasts, a consistent picture emerges: temporal dependence in subjective and objective measures is largely insensitive to motor factors and simple response bias, but is shaped by elapsed time and measurement sensitivity, which modulate the decay constant and noise level revealed in our simulations. When sensitivity is high, both subjective and objective measures express an underlying exponential autocorrelation; when sensitivity is lower, the same process can appear approximately linear, especially for objective performance. These findings reinforce the view that long-range dependencies in subjective reports reflect a slowly evolving latent state rather than being reducible to simple response repetition. A slowly varying perceptual threshold provides one candidate implementation of this state.

### Subjective and objective reports share a component of their temporal fluctuations

A remaining question is whether temporal dependence in subjective reports reflects a slowly varying process shared with objective performance, or an independent drift in subjective response criterion^60–63^. To distinguish these possibilities, we quantified trial-by-trial subjective–objective coupling, that is, the correspondence between subjective reports and objective accuracy, and examined its temporal autocorrelation. In all simulations, subjective responses are transformed deterministically from its threshold, whereas objective responses vary probabilistically (Fig. 6).

**Fig. 6.**
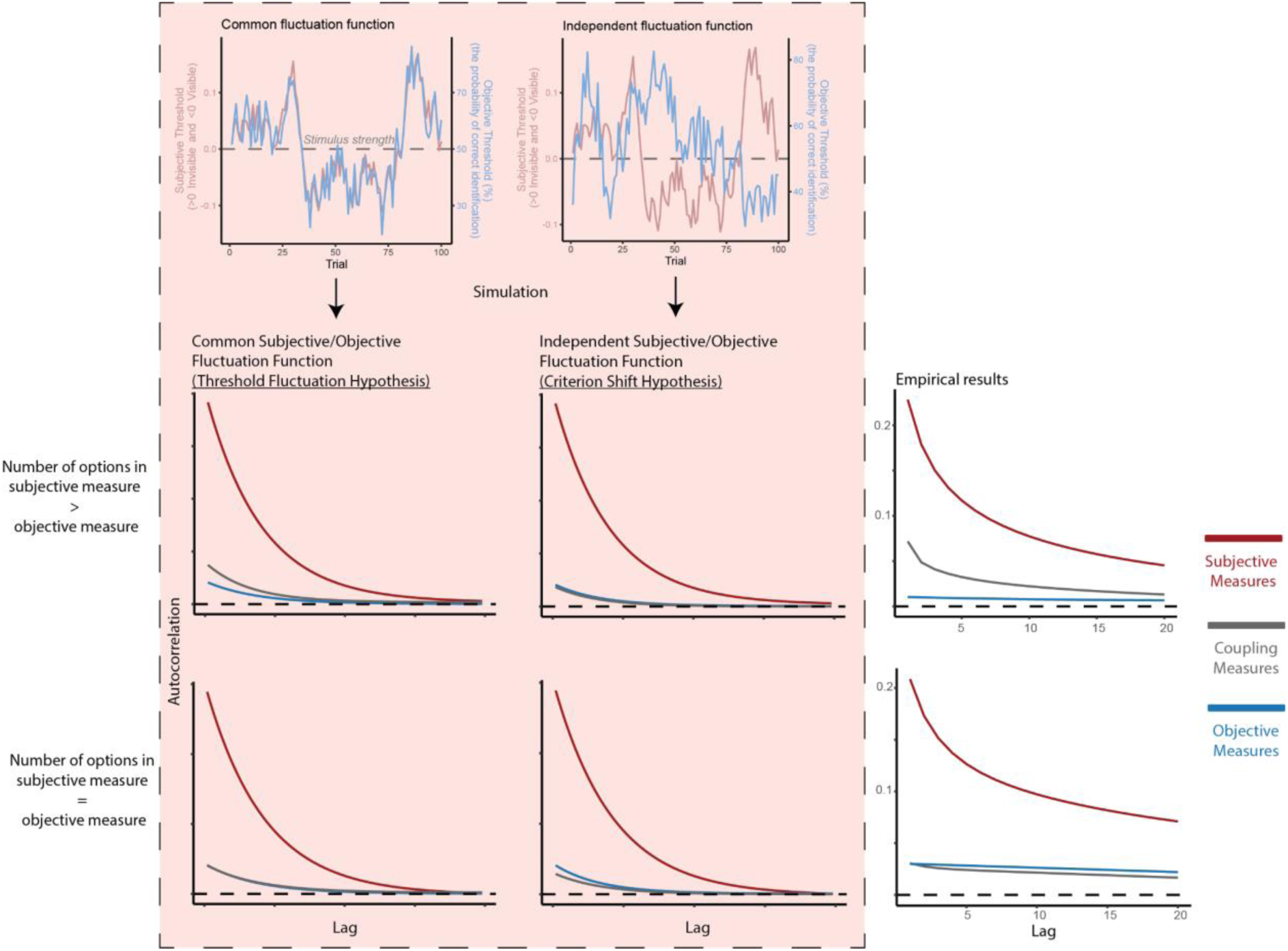
Simulated trial-wise fluctuations and empirical autocorrelation functions dissociate common threshold fluctuation from criterion shift. Top, schematic examples of the latent trial-by-trial processes used in the simulations. Subjective responses were generated from a deterministic transformation from the subjective threshold (e.g., all responses are visible when threshold is below stimulus strength (i.e., 0)), whereas objective responses originated from a probabilistic transformation from the objective threshold (e.g., the further the threshold is below stimulus strength (i.e., 0), the more likely response is correct). Under the common fluctuation function (left), subjective and objective measures inherit temporal structure from a shared source. Under the independent fluctuation function (right), subjective and objective thresholds evolve separately. Bottom, predicted autocorrelation functions for subjective, objective, and their coupling measures under these two hypotheses, shown for cases in which the subjective measure has more response options than the objective measure (upper row of lower panels) or the same number of response options (lower row of lower panels). Right, corresponding empirical autocorrelation functions. Red, subjective measures; grey, subjective-objective coupling measures; blue, objective measures. Two diagnostic contrasts are critical. First, when the subjective measure has more response options than the objective measure, independent source hypothesis predicts that subjective-objective coupling autocorrelation aligns closely with the objective autocorrelation; thus, the extent to which the subjective-objective coupling curve rises above the objective curve indexes the contribution of a common threshold fluctuation. Second, when subjective and objective measures have the same number of response options, any residual criterion shift predicts that the subjective-objective coupling autocorrelation should fall below the objective autocorrelation; thus, the extent to which the subjective-objective coupling curve falls below the objective curve indexes the contribution of the criterion-shift account.

We simulated two alternatives. In the shared-source model, subjective and objective responses inherited temporal structure from a common slowly varying latent process, with independent readout noise. In the independent-source model, the two response streams were generated from independent latent processes, as expected if subjective temporal dependence primarily reflected criterion drift. The models made distinct predictions for the position of the coupling measure relative to subjective and objective measures, depending on whether subjective and objective measures had unequal or equal numbers of response options.

The empirical data showed signatures of both processes. When subjective measures had more response options than objective measures, coupling autocorrelation lay above objective autocorrelation but below subjective autocorrelation (Fig. 6), consistent with a temporal component shared between subjective and objective reports. When the two measures had equal numbers of response options, coupling autocorrelation fell slightly below objective autocorrelation, consistent with an additional subjective-specific component.

Thus, temporal dependence in subjective reports cannot be explained solely by an autonomous drift in reporting criterion. Instead, the results are consistent with a slowly evolving latent process shared between subjective and objective reports, together with an additional subjective-specific component, potentially reflecting criterion drift or confidence carryover. The present analysis does not identify the shared latent variable itself, although fluctuations in perceptual threshold provide one candidate mechanism.

### DLPFC perturbation modulates the temporal dependence of subjective confidence

Having established that subjective reports contain a slowly evolving temporal component, we next asked whether its timescale could constrain its neural implementation. This correspondence does not localize the source of the behavioral dynamics, but it motivates asking whether perturbation of a higher-order cortical region can alter their expression in subjective reports. We therefore converted the exponential decay constants to physical time for 14 datasets in which trial duration was available (Fig. 7a). Across datasets, characteristic timescales (that is the inverse of decay constant) generally exceeded 5 s, while autocorrelation functions approached their asymptotes over approximately 30–100 s (Supplementary Fig. S2). These behavioral timescales are considerably slower than the intrinsic timescales typically reported in the sensory cortex and overlap with the longer timescales observed in association and prefrontal cortex^41,42,45–48^. Although this correspondence cannot localize the underlying process, it identifies higher-order cortical regions with long intrinsic timescales as candidate neural substrates.

**Fig. 7.**
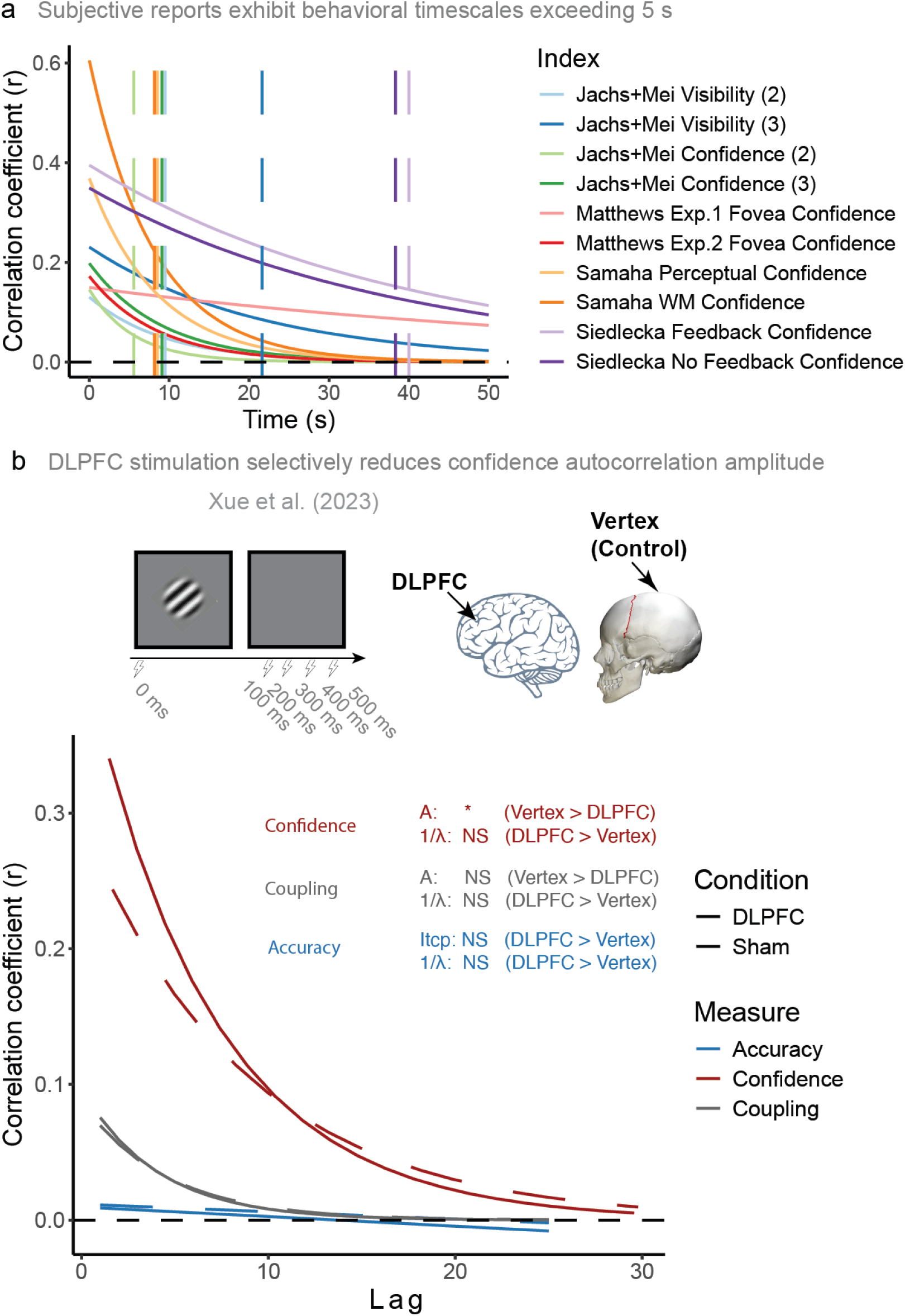
Long behavioral timescales of subjective perception and their modulation by DLPFC stimulation. **a,** Exponential fits of autocorrelation functions for subjective visibility and confidence, replotted in physical time for datasets with documented trial durations. Colored curves show the fitted decay functions for each dataset, and the vertical dashed lines indicate the corresponding characteristic timescales, defined as the inverse of λ. Across 14 datasets, timescales generally exceeded 5 s and in several cases extended into the tens of seconds, consistent with the long intrinsic timescales reported for the prefrontal cortex. **b,** Top, schematic of the paradigm from Xue et al.^64^, in which single-pulse TMS was delivered to right dorsolateral prefrontal cortex (DLPFC) or a vertex (control) condition while participants performed a visual discrimination task and reported confidence. Bottom, fitted autocorrelation functions for confidence, subjective-objective coupling and objective performance. Relative to vertex stimulation, DLPFC stimulation significantly reduced the amplitude of confidence autocorrelation, without reliably changing its decay constant. No significant effects were detected for subjective–objective coupling or objective performance, suggesting a role of DLPFC in the readout of slowly fluctuating subjective thresholds.

We therefore asked whether perturbing prefrontal cortex alters this temporal signature. We reanalysed data from Xue et al.^64^, in which single-pulse TMS was delivered to right dorsolateral prefrontal cortex (DLPFC) or vertex (control site) at 0, 200, 300, 400 or 500 ms after stimulus onset, while participants discriminated the orientation of a noisy Gabor patch and reported confidence (Fig. 7b). Our simulations provided a specific prediction: if DLPFC contributes to the expression or reading out the slow fluctuations expressed in confidence, perturbation should reduce their autocorrelation amplitude, whereas their characteristic decay rate need not change (Fig. 4g).

This is what we observed. Relative to vertex stimulation, right DLPFC stimulation significantly reduced the amplitude of the confidence autocorrelation (A: *β̂* = −0.129, SE = 0.050, P = 0.010), while leaving the decay constant unchanged (λ: *β̂* = −0.033, SE = 0.034, P = 0.323; Fig. 7b). By contrast, objective performance exhibited no detectable TMS effect in the linear fit (intercept: *β̂* = 0.002, SE = 0.006, P = 0.735; slope: *β̂* = 0.0001, SE = 0.0003, P = 0.589; Fig. 7c). The subjective–objective coupling measure showed numerically lower amplitude following DLPFC stimulation, but neither this effect nor the change in decay constant was significant (A: *β̂* = −0.010, SE = 0.024, P = 0.681; λ: *β̂* = −0.033, SE = 0.068, P = 0.623).

Together, the long intrinsic timescales of subjective measures and the reduction of confidence-autocorrelation amplitude following DLPFC stimulation identify the prefrontal cortex as one candidate contributor to the expression or readout of these slow subjective dynamics. The absence of detectable effects on the decay constant, objective performance, and subjective–objective coupling leaves open whether DLPFC participates in the underlying slow latent process itself or instead influences how that process is translated into subjective confidence. The principal contribution is identifying a reproducible temporal property of subjective perception that future neural and computational accounts of consciousness must explain.

## Discussion

Conscious perception is commonly approached as a series of independent snapshots: given the same sensory input, why is a stimulus consciously perceived on one trial but not another? Our findings suggest that contrary to what is commonly assumed, there are systematic temporal dependencies, such that the probability of conscious perception itself carries a history. Across diverse datasets, subjective visibility and confidence were not independent from one moment to the next but exhibited temporal dependencies spanning multiple trials and several seconds. These dependencies were strongest and most consistently exponential for subjective reports, whereas objective performance showed a substantially weaker, approximately linear profile. Conscious access therefore appears to depend not only on the sensory evidence available at the present moment, but also on a slowly evolving internal state inherited from preceding perceptual events.

Thus, our results indicate that the conditions governing whether sensory information becomes subjectively available evolve slowly over time. Several observations argue against this structure arising simply from response repetition or motor perseveration. Interspersing liminal trials with multiple supra- or subliminal trials eliciting predominantly visible or invisible responses did not abolish temporal dependence between liminal trials. Moreover, experiments with markedly different trial durations converged when timescales were expressed in physical rather than trial time, and raw identification responses showed only weak autocorrelation (Supplementary Fig. S3). Thus, the relevant temporal memory appears to reside not simply in the preceding response but in an internal state evolving with elapsed time. Such dynamics are largely invisible to conventional analyses that average across trials and implicitly treat successive observations as independent samples from a stationary perceptual process. Autocorrelation provides a complementary window onto this latent temporal context.

What, then, is fluctuating? One possibility is that the temporal dependence arises entirely downstream of perception, because participants slowly shift the criterion used to report a stimulus as visible or their decision as confident^60–63^. Our results argue against this being the sole explanation. Objective performance also exhibited reliable, although weaker, temporal dependence. Moreover, trial-by-trial feedback provides an informative manipulation. In Siedlecka et al.^50^, objective performance was approximately linear without feedback but became better described by an exponential function when performance feedback was provided. Our simulations show that increasing readout noise can make an underlying exponential process appear shallow and approximately linear. Thus, the different temporal profiles of subjective and objective measures need not imply completely independent processes; they may partly reflect a common slowly varying state expressed through readouts of different sensitivity.

The analysis of subjective–objective coupling further supports this interpretation. Empirically, its temporal profile more closely resembled predictions from simulations in which subjective and objective responses inherited temporal structure from a common latent process than those in which they fluctuated independently. Yet the data also contained evidence for an additional subjective-specific component. This provides a natural connection to confidence leak, whereby confidence on one judgment biases subsequent confidence judgments, even across tasks or domains^14,32–34^. Rather than viewing confidence leak and latent perceptual fluctuations as competing explanations, our findings suggest at least two components: a relatively local carryover in subjective criterion and a slower latent fluctuation that also leaves a weaker signature in objective performance. The latter is compatible with fluctuations in perceptual threshold, although the present analyses do not uniquely identify the underlying variables, which might covary with fatigue, alertness, attention etc.

Although subjective and objective responses share a common underlying latent fluctuating process, we found distinct autocorrelation profiles. The simulation in Fig. 6 shows that the much weaker autocorrelation strength of objective accuracy might result from its probabilistic transformation from its threshold. This probabilistic transformation involves more noise in the read-out process than the deterministic transformation of the subjective threshold, which is consistent with the results in Fig. 4g, where more noise suppresses the strong exponential function to a weak linear-like one. Therefore, the current study identifies two factors that dissociate subjective and objective measures of consciousness (i.e., relative blindsight). One is the presence of independent criterion shifts exclusive to each respective measure. The other is the stronger read-out noise for objective responses from its latent threshold.

Another important boundary condition emerged in paradigms involving divided attention and spatially distributed or peripheral stimuli. Temporal dependence in subjective measures was markedly reduced when stimuli were presented under divided attention^15,65–67^. We considered two non-exclusive explanations. One is that attention modulates the strength of the underlying fluctuation, such that focused attention on a single foveal location allows the exponential structure to fully manifest, whereas divided or unfocused attention attenuates it. Under this view, strong autocorrelation should re-emerge whenever covert attention is consistently directed to the target stimulus a priori, even at varying retinotopic locations. This hypothesis aligns well with the global neuronal workspace theory of consciousness^68,69^. This theory suggests that consciousness depends on sufficient attention and stimulus strength^65^. A second possibility is that threshold fluctuations are themselves retinotopically organized, with partially independent processes at different positions in visual space. In that case, switching the target location across trials would disrupt autocorrelation even under focused attention. This possibility would be compatible with previous results demonstrating that perceptual learning alters the threshold of objective performance in a retinotopically specific manner, consistent with receptive fields of visual areas up to V4, while threshold changes for subjective visibility likely arose from higher areas^28^. Distinguishing these hypotheses will require future experiments that systematically manipulate attention and location while tracking temporal dependence within and across positions.

A final contribution of our findings is the introduction of a temporal constraint on candidate neural correlates of conscious access. Classical NCC approaches seek neural activity that distinguishes seen from unseen stimuli at a given moment^44,70^. Our results suggest that candidate mechanisms should additionally reproduce the timescale over which the probability of conscious access fluctuates. Such activity would not necessarily represent the content of the current experience, but rather the momentary capacity for incoming sensory information to become consciously accessible. The correspondence with long cortical timescales, together with the selective effect of DLPFC stimulation on confidence autocorrelation, makes prefrontal and association networks plausible candidates, while slow fluctuations in alpha power and aperiodic infraslow activity provide candidate neural signals^23,24,27,71^. Directly relating these neural fluctuations to behavioral autocorrelation will provide a critical test of this proposal.

Taken together, our results show that conscious perception is shaped by slow temporal dynamics that leave a robust signature in subjective awareness. These dynamics are consistent with a slowly evolving latent state, for which fluctuations in perceptual threshold provide one candidate implementation. The DLPFC findings further identify the prefrontal cortex as one possible contributor to the expression or readout of these dynamics, rather than uniquely establishing their neural source. These observations introduce a new temporal constraint on theories and neural candidates of consciousness: candidate mechanisms should explain not only why a stimulus is seen or unseen at a given moment, but why the probability of seeing it changes over time. Future work should move beyond trial-averaged contrasts and test for how intrinsic neural timescales, attention, retinotopic organization, and cross-domain generalization jointly shape our momentary capacity for conscious access.

## Methods

This study was conducted based on the re-analysis of existing data. To test the autocorrelation function of subjective and objective measures of consciousness, we set three criteria for dataset selection: 1. stimuli with the same physical characteristics presented on the liminal threshold multiple times, which controls the physical input across trials; 2. at least one subjective detection task (visibility/confidence ratings) and one objective discrimination task on the same stimulus to obtain both subjective and objective measures on every trial; 3. equal number of liminal trials in each block/run to enable a standard estimate of lag within a block. The datasets were not identified through a systematic literature search and therefore should not be regarded as an exhaustive meta-analytic sample. Rather, we used this heterogeneous set of available datasets to evaluate the reproducibility and generality of the temporal dependencies across different experimental contexts. Where estimates were quantitatively pooled across datasets, we refer to this procedure as a cross-dataset random-effects synthesis. Re-analysis of only a subset of available datasets in the literature to the author’s knowledge is conducted that meet the aforementioned requirements.

### Subjects and Design

20 existing datasets reported in 9 articles are reanalyzed in this study. The raw dataset from 5 articles are obtained from the link (https://osf.io/s46pr/) reported in the confidence database^56^. A brief introduction to the subjects and design of each study is provided below. We recommend accessing the original articles for detailed descriptions. A tabular summary of the dataset and experimental parameters can be seen in **Supplementary Table 4**.

#### Zheng et al.^8^

81 participants from Zhejiang University were recruited using monetary compensation. Only the participants whose visibility on the liminal trials is lower than 80% and higher than 20% are selected for analysis, which leads to 59 valid participant datasets.

On each trial, participants performed a Stroop priming task under continuous flash suppression. A 500-ms prime word (meaning either yellow or blue) in Chinese was presented to the non-dominant eye, whereas a continuously flashing mask was displayed to the dominant eye. This prime is followed by a 200-ms target color block (either in yellow or blue) that is presented to both eyes. Participants were instructed to report the color of the target as soon as possible as Task 1. Upon response to T1, they are required to report the visibility and identification of the prime in a prime assessment grid. Three levels of prime visibility (supraliminal, liminal, and subliminal) are determined by the prime contrast (high, medium, and low). In the two baseline blocks, 94 out of 100 trials are liminal trials. In the two visible context blocks, 20 are liminal trials and 74 are supraliminal trials, which is similar to two invisible context blocks (20 liminal trials and 74 supraliminal trials). There are 6 catch trials in every block type.

#### Matthews et al.^54^

Eight participants from Monash University were recruited for Experiment 1 and thirteen for Experiment 2. Only the data of two-response dual-task conditions in Experiment 1 and 2 are selected in this analysis, as the stimulus onset asynchronies (SOAs) of the central and peripheral stimuli were not fixed within a block in the other conditions. Participants with lower than 1.6 or larger than 3.4 average confidence rating in the four-level confidence rating scale on the liminal trials and incomplete participation were excluded from the analysis.

There were two tasks on two stimuli on each trial: central letter discrimination task for foveal stimuli and peripheral discrimination task for peripheral stimuli. The foveal stimuli consist of five letters all in random angle (one at fixation and four stimuli 3° from fixation). They are either all the same (all ‘T’ or ‘L’) or contain one single differing letter (one ‘T’ in four ‘L’). After a fixed delay, five ‘F’ appeared at the same five locations to mask the letters that appeared before. A peripheral stimulus (2.5°) was displayed at one of the four corners of the screen (8° × 10°). In Experiment 1, a face stimulus was followed by an achromatic mask. In Experiment 2, a color disk was followed by a chromatic mask. Identifications of each stimulus (2 alternatives) and subjective confidence (1-4 levels) are jointly reported with a single mouse click on the assessment grid.

#### Del Pin et al.^55^

Twenty-two and nineteen participants were collected for Experiment 1 and 2, respectively. The participants who reported lower than 1.6 or larger than 3.4 average visibility on a PAS scale ranging from 1 to 4 were excluded from data analysis. This leads to the exclusion of three participants in Experiment 1.

A 1000-ms blank screen with fixation cross precedes the 250-ms stimulus frame. This frame contains eight objects (2.41° × 2.41°) laid out in an imaginary circle (4.82° × 4.82°) around the fixation cross. Every object was placed in a black square. Afterwards, a 100-ms blank screen is followed by a red line pointing towards one of the eight empty black squares for 500 ms. After a 1400 ms blank screen delay, participants were required to report the identity of the object at the cued location in a 2AFC task (displayed in object picture or words) and visibility of the object on the PAS scale, respectively. The experiment consists of 200 trials with a regular break after every 25 trials. Half trials used an object picture in the object identification task, and the other half used words.

#### Samaha & Postle^49^

Twenty participants from University of Wisconsin-Madison participated in the Experiment 1 of this study. Experiment 2 does not apply to this analysis as the delay duration varies across trials. The same data exclusion was applied as Matthews et al.^54^, which excluded 4 participants from this experiment.

This experiment contains two conditions: perceptual condition and working memory (WM) condition. The perceptual task required participants to estimate the orientation of a target grating, briefly presented in noise, by adjusting a visible probe grating’s orientation and rating their confidence on a scale from 1 to 4. The VSTM task is identical to the perceptual task except including a fixed delay (7s) between the target and probe. Perceptual and VSTM tasks were conducted in separate blocks, totaling 240 perceptual and 180 VSTM trials per subject.

#### Siedlecka et al.^50^

Thirty-seven healthy volunteers with normal or corrected vision participated in the experiment, receiving a small payment. The data of six participants was excluded from the analysis driven by the same criteria as datasets above.

The experiment, conducted on PCs using PsychoPy, utilized LCD monitors (1920 × 1080 resolution, 60 Hz). Stimuli were circular gratings in white noise, oriented left or right, calibrated for each participant. The 4-level PAS scale, shown after orientation responses, ranged from “no experience” to “a clear experience.” The procedure included 15 training trials with feedback, a calibration session for contrast adjustment, followed by a break. Participants then completed two experimental blocks (with and without feedback), each containing 200 trials. Trials began with a blank screen (500 ms), a fixation cross (500 ms), and a grating (33 ms). Participants identified the grating orientation and rated their experience on the PAS. The feedback condition included accuracy feedback after PAS ratings. The total response time was limited to 3 seconds.

#### Jachs et al.^52^

Thirteen, eighteen, and ten healthy volunteers participated in Experiment 1, Experiment 2, and Experiment 3, respectively. Three, five, and zero participants from three experiments reported visible experience on less than 20% or more than 80% trials, respectively. Therefore, they were excluded from the analysis.

In Experiment 1, each trial started with a fixation display (500 ms) followed by a 2AFC task to determine the orientation of a Gabor patch (3 cm diameter) presented for 35 ms, followed by a backward mask (353 ms). They reported awareness, orientation, and confidence on each trial, completing 10 blocks of 50 trials. Participants underwent training and stimulus calibration phases, followed by 30 training trials using the calibrated luminance. Similar to Experiment 1, Experiment 2 used a perceptual awareness scale (1-3 rating) for a more detailed measure of awareness. Similar to previous experiments, participants reported awareness and confidence on a 1-2 scale and completed eight blocks of 50 trials each in Experiment 3.

#### Mei et al.^53^

Fifteen participants participated for monetary compensation in Experiment 1, and sixteen in Experiment 2. One participant from each experiment was excluded prior to analysis as they reported visible experience on less than 20% of the liminal trials.

In Experiment 1, each trial involved participants first reporting their prospective belief of success (low or high) for the upcoming task. A Gabor patch, oriented 40 degrees left or right, was presented for 35 ms at the threshold of visual awareness, followed by a 353 ms mask. Participants then rated their awareness of the Gabor, identified its orientation, and rated their confidence in their response. Each participant completed 12 blocks of 50 trials (600 total), with breaks between blocks. Participants reported awareness, orientation, and confidence without a response deadline. Experiment 2 is completely identical to Experiment 1 except that participants need to report their decision to engage attention to task (high or low).

#### Sand & Nilsson^51^

Sixty-seven participants were recruited. We excluded participants who reported visible on less than 20% trials or more than 80% trials. This leads to thirty-one participants valid for the analysis.

Every trial starts with a fixation cross (500-900 ms) and is followed by a 6-ms prime word (either RED or BLUE) backward masked by a 140-ms mask which consists of 6 # symbols. Prime and mask are separated by a blank screen as interstimulus interval (ISI). A 140-ms rectangle in blue or red, i.e., the target, follows the mask offset. Speeded response to target color was required since target onset, and visibility (“No Percept”, “Unclear Percept”, and “Clear Percept”) and identification (“RED” or “BLUE”) of the prime were to be reported on a prime-assessment grid using a single mouse click. 330 trials in total consist of 240 liminal trials (ISI = 25 ms), 60 high visibility trials (ISI = 100 ms), and 30 no-prime catch trials.

#### Xue et al.^64^

Seventy-two participants were recruited. The data of sixty-two participants are publicly available (forty-three in DLPFC and nineteen in Vertex control groups). Among these, eight from the DLPFC group and five from Vertex control group were excluded who reported lower than 1.6 or larger than 3.4 average confidence rating on a scale ranging from 1 to 4.

On each trial, participants judged the orientation (left vs right) of a noisy Gabor patch presented for 100 ms and simultaneously reported their confidence on a 4-point scale via a single key press. After training and a staircase procedure to set individual contrast thresholds, they completed four runs of 5 × 40-trial blocks (800 trials total). Single-pulse TMS was delivered on every trial either to right DLPFC or to vertex (between-subjects factor) at 0, 200, 300, 400, or 500 ms after stimulus onset, with the five delays intermixed in pseudorandom order.

### Data Analysis

#### Modelling the autocorrelation functions of subjective, objective measures, and subjective-objective coupling

No transformation is performed on visibility, confidence, or accuracy measures. Subjective-objective coupling is calculated to represent the consistency between subjective measures (visibility and confidence) and objective measure (accuracy). Both subjective and objective measures are linearly transformed to a scale between 0 (invisible, no confidence, or maximally incorrect) and 1 (fully visible, fully confident, or completely correct) on a trial-by-trial basis, regardless of the number of possible responses. Subjective-objective coupling is computed as one minus the absolute distance between the linearly transformed subjective and objective measure, which also ranges between 0 (fully inconsistent) and 1 (fully consistent).

This autocorrelation analysis is only performed on the liminal trials in each block. A pairwise correlation between all possible pairs of a measure (visibility, confidence, or accuracy) given a specific lag within a run was conducted for each participant. This computation was applied to all three measures for each experiment. Here, the lag refers to the distance between liminal trials only (see Fig. 1 and Fig. 2). We computed this autocorrelation coefficient from lag 1 trial to lag N trials for each participant. N is the 60% of the total number of liminal trials in a block. This leads to a data frame that contains the correlation coefficient per lag per participant per dataset. We also calculated the standardized lag by multiplying the lag with the inverse of the proportion of liminal trials in each block. This standardized lag denotes the average lag between trials (including liminal and all other trial types) that underlies each autocorrelation coefficient (see Fig. 3). This serves to enable fair comparisons between datasets that have different proportions of liminal trials in a block.

We conducted nonlinear least squares fitting to fit the autocorrelation data of all subjects with the exponential decay function using the *gsl_nls* function in R package *gslnls*^72^. The mathematical formula of the exponential decay function implemented in the *gsl_nls* function is followed:

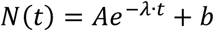

Three parameters are to be estimated in the function: A, λ, and b. As shown in Fig. 1, A refers to the amplitude of the function, i.e., the lag between N(0) and N(∞). λ represents the exponential constant, i.e., determining how fast the exponential function decays. The inverse of the exponential constant is the timescale, denoting the lag at which the function decays to 36.8%, i.e., the inverse of the mathematical constant *e*, of the amplitude. b stands for the offset, i.e., N(∞). The independent variable t is the standardized trial-to-trial lag.

We also applied a linear function to fit the autocorrelation coefficients of three measures as a function of standardized lag using *lm* function in R package *stats*. The slope and intercept of the linear regression are estimated.

We employed likelihood ratio and Wald-Type tests to test which of the two mathematical functions explain more variance in the autocorrelation data. We used *anova* function from R package *metafor*^63^ for this analysis.

To complement these nested-model tests, we additionally compared the exponential and linear models using information criteria, which accommodate the comparison of non-nested models and do not rely on the asymptotic reference distribution of a likelihood-ratio statistic. This is relevant here because the linear model is not a conventional parameter restriction of the exponential model but rather approximates its behaviour in the limit of a small decay constant (λ → 0), placing the implied null on the boundary of the parameter space. For each dataset and measure, we computed the small-sample-corrected AIC (AICc) for both the exponential decay model and the linear model. AICc was computed as AIC + 2k(k + 1)/(n − k − 1), where k is the number of estimated parameters (including the residual variance; k = 4 for the exponential model and k = 3 for the linear model) and n is the number of observations; AICc was preferred given the modest number of observations in several datasets. Log-likelihoods were obtained under a Gaussian error model and computed on the identical set of autocorrelation coefficients used for both fits, ensuring that the criteria were directly comparable across the two functional forms. For each comparison we report Akaike weights derived from AICc as an interpretable measure of the relative support for each model. Because autocorrelation coefficients are not independent across lags, we report differences in information criteria and model weights rather than interpreting the absolute values of the criteria. Full per-dataset criteria and weights are provided in Supplementary Table 1.

We followed the same protocol for the analysis on every dataset.

To test whether each estimated parameter is significantly different from zero, we conducted meta-analysis on the three parameters in the exponential decay function for visibility and confidence data and two parameters in the linear function for accuracy data.

We obtained the estimate and standard error of each parameter from the output of the functions reported above corresponding to each dataset. We applied the generic inverse variance meta-analysis using the *metagen* function in the R package *meta*^73^. Between-study variance is estimated using restricted maximum-likelihood estimator (REML)^74^. As all studies employ different experimental paradigm and data is collected from different populations, the random effect model is applied. We adopted a method from Hartung and Knapp^75,76^ to estimate the confidence interval for random effect estimates. The forest plot was generated using the *forest* function from R package *meta*.

To understand the effect of specific experimental parameters on the autocorrelation functions, we contrasted the autocorrelation functions between different datasets. We added a new variable called *condition* in the nonlinear least square fitting function for the contrast of each experimental parameter. We contrast-coded two datasets with -0.5 and 0.5, respectively. Here, the term *condition*, simplified as *cond*, refers to the only experimental parameter that differs across the two contrasted datasets. To test whether this experimental parameter significantly influences the three parameters of the exponential decay function, we implemented the estimation of the following function using the *gsl_nls* function.

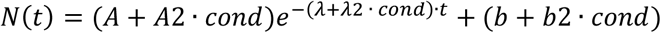

Three more parameters are to be estimated in this equation, including A2, λ2, and b2. They are the coefficients of the condition variable. If A2, λ2, or b2 are significantly different from zero, there is significant difference in the amplitude, exponential constant, or offset between the autocorrelation functions of the two contrasted datasets. The statistical inferences on the differences in the parameters of each contrast are noted in Fig. 3 and Fig. 7b.

#### Testing two hypotheses of threshold fluctuation function through simulation

In the previous analysis, we computed how subjective visibility and objective accuracy changes in time despite constant strength of stimulus input. If we define subjective and objective perceptual threshold as the minimal stimulus strength that produces subjective visible experience and objectively correct identification, there must be a fluctuating subjective and objective threshold to account for the changes in visibility and accuracy responses. To understand what is the underlying threshold fluctuation function that leads to the exponential decay function in the autocorrelation function of subjective measures and linear function in the autocorrelation function of objective measure, we simulated two hypotheses of threshold fluctuation function: periodic fluctuation and aperiodic fluctuation with temporal dependency. Our simulations of all three hypotheses assume 50 participants, 2 blocks per participant, and 100 trials per block.

The periodic fluctuation hypothesis suggests that perceptual threshold fluctuates periodically in time based on the following formula (Fig. 4d):

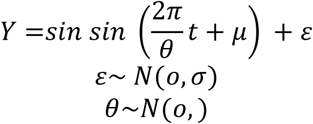

Here, θ is the period of the periodic function of the hypothetical individual. μ is the phase. ε stands for the noise that overlays the periodic function. σ is the standard deviation of the noise. ν is the standard deviation of fluctuation periods across individuals. An example of the threshold fluctuation can be seen in Fig. 4e. Similar to the previous hypothesis, we computed its autocorrelation function after transforming the threshold to binary data format. As shown in Fig. 4f, this hypothesis also predicts periodicity approaching zero in its autocorrelation function. The period in the autocorrelation function depends on the average period of the threshold fluctuation function across participants.

The aperiodic fluctuation hypothesis suggests that perceptual threshold fluctuates aperiodically with temporal dependency. In the simulation, we first set the average threshold of a hypothetical participant by drawing a random number μ out of a normal distribution (mean = 0; sd = ν). We then generated an array of random numbers (X_i_) that is uniformly distributed between -1 + μ and 1 + μ. We then computed the weighted sum of past values X_i_, where more recent values have exponentially higher weights due to the exponential factor *γ^i^*^−*t*+1^. The threshold at timepoint t is determined by the weighted sum of past values in X_i_ divided by a normalizing factor 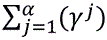 in addition to ε, noise in the response system.

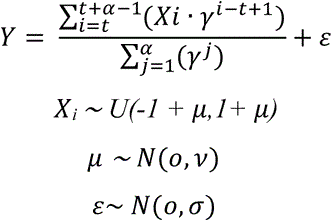

Here, t stands for time. *α* means the length of the temporal dependency window, indicating how many past values of X are considered. γ refers to the base of the exponential weights, controlling how the influence of past values decays over time. *μ* is the mean threshold of a subject, influencing the range of the X_i_ values. ν is the standard deviation of the mean threshold *μ* across subjects, indicating the variability in thresholds. ε is the noise term, representing random fluctuations in the response Y. *σ* is the standard deviation of the noise ε.

This hypothesis predicts an autocorrelation function in the exponential decay pattern (Fig. 4i, j, k). Specifically, we found that the base of exponential weights determine how the exponential constant in the autocorrelation function (Fig. 4i). An exponential decay pattern can be observed when the noise ε is low, whereas the function follows a linear pattern when the noise is high (Fig. 4j). Lastly, the larger the standard deviation of the mean threshold across participants ν is, the more participants’ thresholds are far beyond or below the strength of the sensory input. This also results in higher offset in the autocorrelation function.

#### Simulating shared versus independent threshold fluctuations for the threshold-fluctuation and criterion-shift comparisons

To evaluate whether the exponential autocorrelation of subjective reports could arise from a shared latent fluctuation with objective performance, or whether it would instead persist even when subjective and objective responses were generated from statistically unrelated sources, we ran a simulation with two parallel architectures, one in which subjective and objective measures were derived from a common latent trajectory and one in which they were derived from independent latent trajectories.

##### Generating the latent evidence

We simulated 1,200 independent observers, each providing a run of 100 trials. For each simulated observer, we drew two independent sequences of 200 random numbers, uniformly distributed between −0.99 and 0.99. One sequence served as the common latent evidence stream, from which both the subjective threshold and the “shared-source” objective measure were derived. The second, independent sequence served as the source for the “independent-source” objective measure, used in the criterion-shift comparison. We additionally drew, per observer, a noise sequence of 100 values from a normal distribution (mean 0, SD 0.1), which was added to the subjective threshold to represent noise specific to the readout of the subjective report.

##### Constructing the threshold trajectories

Each latent stream was converted into a 100-sample threshold trajectory using an exponentially weighted moving average with a 100-sample window, in which the most recent samples in the window contributed most strongly and earlier samples contributed progressively less (weights increasing geometrically by a factor of 1.05 per step). Because each successive trial’s window overlaps almost entirely with the previous trial’s window, this procedure imparts genuine serial dependence to the resulting trajectory, producing an autocorrelation function that decays approximately exponentially with lag by construction.

Using this procedure, we generated:

- a subjective threshold trajectory, from the common latent stream plus the subjective-specific noise term described above;
- a shared-source objective threshold trajectory, from the same common latent stream, with independent trial-by-trial noise (SD 0.1) added instead of the shared noise term; and
- an independent-source objective threshold trajectory, from the second, unrelated latent stream, with its own independent trial-by-trial noise (SD 0.1).

##### Deriving binary and graded responses

The subjective threshold trajectory was converted into a binary response (visible/invisible) by thresholding at zero for binary subjective responses (the number of options for subjective measure equates the number of options for objective measure), or separately into a four-level graded response by dividing the trajectory into four ordered bins around zero for four level subjective responses (the number of options for subjective measure outnumbers the number of options for objective measure). Each objective threshold trajectory was converted into a graded probability of a correct response: values below zero were mapped onto a probability of being correct that increased linearly as the threshold value became more negative (scaled so that the probability reached 1 at a specific threshold value), while values at or above zero were assigned a floor probability of 0.5, corresponding to chance-level performance, assuming no blindsight response. A binary correct/incorrect outcome was then sampled on every trial according to this probability, separately for the shared-source and the independent-source objective trajectories.

##### Deriving the subjective-objective-coupling measure

For each of the two objective trajectories (shared-source and independent-source), we computed subjective-objective coupling, i.e., a measure of trial-by-trial coupling between the subjective and objective responses, separately for two response-format combinations:

- a matched-format version, in which the binary subjective response (0 or 1) was compared directly to the binary objective outcome (0 or 1), computed as one minus the absolute difference between the two: agreement (1) or disagreement (0);
- an unmatched-format version, in which the four-level subjective response was rescaled to the 0, 0.33, 0.66, or 1 and compared to the binary objective accuracy (0 or 1), with the coupling measure defined as one minus the absolute difference between the two. Similarly, 1 stands for maximal agreement, whereas 0 minimal agreement.

This yielded four coupling curves in total: matched- and unmatched-format coupling for the shared-source objective measure, and matched- and unmatched-format coupling for the independent-source objective measure.

We followed the same algorithm as mentioned above to compute autocorrelation as a function of lag and fit the exponential decay function of subjective, objective, and subjective-objective responses.

##### Assembling the four comparison plots

Each of the four plots displays three fitted decay curves on a common lag axis:

- Threshold-fluctuation (common source), subjective measure options > objective measure options: the four-level subjective response, the shared-source objective response, and their unmatched-format coupling measure.
- Threshold-fluctuation (common source), subjective measure options = objective measure options: the binary subjective response, the shared-source objective response, and their matched-format coupling measure.
- Criterion-shift (independent source), subjective measure options > objective measure options: the four-level subjective response, the independent-source objective response, and their unmatched-format coupling measure.
- Criterion-shift (independent source), subjective measure options = objective measure options: the binary subjective response, the independent-source objective response, and their matched-format coupling measure.

The rationale for this contrast is that, in the threshold-fluctuation plots, the objective measure is generated from the same latent trajectory as the subjective threshold, so any autocorrelation shared between the subjective and coupling curves reflects a genuine common fluctuation process. In the criterion-shift plots, the objective measure is generated from a statistically independent latent trajectory, so any residual autocorrelation observed in the coupling curve under this architecture cannot be attributed to a shared perceptual threshold, and instead indexes the kind of spurious dependence that a criterion-shift or response-carryover account would predict.

## Author contributions

Conceptualization: D.T., L.M. and Z.Z.; Investigation: D.T., L.M. and Z.Z.; Supervision: D.T., Y.C., and L.M.; Visualization: Z.Z.; Writing - original draft: Z.Z.; Writing - review & editing: D.T., Y.C., L.M., and Z.Z..

## Acknowledgement

We thank Alex Lepauvre, Qiyuan Zeng, Qian Chu, and other members of the Research Group Neural Circuits, Consciousness, and Cognition at Max Planck Institute for Empirical Aesthetics and Predictive Brain Lab at Ruhr-University Bochum for comments on the different stages of the project.

## Data and code availability

All data and code used in the analysis reported are available under the OSF project: https://doi.org/10.17605/OSF.IO/X7N9Y

## Supplementary Materials

**Table S1.** Statistical summary of model comparison between the exponential decay and linear fit to each dataset of each respective measure.

| Dataset | Measure | n_exp | n_lin | w_exp_AICc | w_lin_AICc | Res.Df | Res.Sum Sq | Df | Sum Sq | F value | Pr(>F) |
| --- | --- | --- | --- | --- | --- | --- | --- | --- | --- | --- | --- |
| Zheng Baseline | Accuracy | 3347 | 3347 | 0.297 | 0.703 | 3345 | 86.142 | -1 | -0.007 | 0.284 | 0.594 |
| Zheng Baseline | Visibility-Accuracy coupling | 3347 | 3347 | 0.324 | 0.676 | 3345 | 84.755 | -1 | -0.013 | 0.532 | 0.466 |
| Zheng Baseline | Visibility | 3347 | 3347 | 1.000 | 0.000 | 3345 | 209.164 | -1 | -4.779 | 78.195 | 0.000 |
| Zheng Vis/Invis | Accuracy | 680 | 680 | 0.267 | 0.733 | 678 | 18.214 | -1 | 0.000 | 0.000 | 1.000 |
| Zheng Vis/Invis | Visibility-Accuracy coupling | 680 | 680 | 0.267 | 0.733 | 678 | 17.451 | -1 | 0.000 | 0.000 | 1.000 |
| Zheng Vis/Invis | Visibility | 680 | 680 | 0.986 | 0.014 | 678 | 40.686 | -1 | -0.626 | 10.582 | 0.001 |
| Jachs Exp.1 | Accuracy | 400 | 400 | 0.273 | 0.727 | 398 | 1.987 | -1 | 0.000 | 0.083 | 0.773 |
| Jachs Exp.1 | Confidence | 400 | 400 | 0.999 | 0.001 | 398 | 1.983 | -1 | -0.080 | 16.800 | 0.000 |
| Jachs Exp.1 | Confidence-Accuracy coupling | 400 | 400 | 0.427 | 0.573 | 398 | 1.645 | -1 | -0.006 | 1.448 | 0.230 |
| Jachs Exp.1 | Visibility-Accuracy coupling | 400 | 400 | 0.332 | 0.668 | 398 | 1.717 | -1 | -0.003 | 0.638 | 0.425 |
| Jachs Exp.1 | Visibility | 400 | 400 | 0.992 | 0.008 | 398 | 2.360 | -1 | -0.068 | 11.720 | 0.001 |
| Jachs Exp.2 | Accuracy | 520 | 520 | 0.715 | 0.285 | 518 | 1.978 | -1 | -0.015 | 3.863 | 0.050 |
| Jachs<br>Exp.2 | Confidence | 520 | 520 | 1.000 | 0.000 | 518 | 3.333 | -1 | -0.327 | 56.272 | 0.000 |
| Jachs<br>Exp.2 | Confidence<br>-Accuracy<br>coupling | 520 | 520 | 0.897 | 0.103 | 518 | 2.091 | -1 | -0.025 | 6.360 | 0.012 |
| Jachs<br>Exp.2 | Visibility-Accuracy<br>coupling | 520 | 520 | 0.266 | 0.734 | 518 | 2.086 | -1 | 0.000 | -0.003 | 1.000 |
| Jachs<br>Exp.2 | Visibility | 520 | 520 | 0.998 | 0.002 | 518 | 5.704 | -1 | -0.157 | 14.661 | 0.000 |
| Jachs<br>Exp.3 | Accuracy | 400 | 400 | 0.288 | 0.712 | 398 | 2.108 | -1 | -0.001 | 0.230 | 0.632 |
| Jachs<br>Exp.3 | Confidence | 400 | 400 | 1.000 | 0.000 | 398 | 2.759 | -1 | -0.207 | 32.261 | 0.000 |
| Jachs<br>Exp.3 | Confidence<br>-Accuracy<br>coupling | 400 | 400 | 0.265 | 0.735 | 398 | 2.292 | -1 | 0.000 | 0.000 | 1.000 |
| Jachs<br>Exp.3 | Visibility-Accuracy<br>coupling | 400 | 400 | 0.460 | 0.540 | 398 | 3.696 | -1 | -0.016 | 1.713 | 0.191 |
| Jachs<br>Exp.3 | Visibility | 400 | 400 | 1.000 | 0.000 | 398 | 5.447 | -1 | -0.344 | 26.794 | 0.000 |
| Matthews<br>Exp.1<br>Fovea | Accuracy1 | 252 | 252 | 0.263 | 0.737 | 250 | 1.523 | -1 | 0.000 | 0.000 | 0.991 |
| Matthews<br>Exp.1<br>Periphery | Accuracy2 | 252 | 252 | 0.263 | 0.737 | 250 | 1.330 | -1 | 0.000 | 0.000 | 1.000 |
| Matthews<br>Exp.1<br>Fovea | Confidence<br>1 | 252 | 252 | 0.775 | 0.225 | 250 | 3.988 | -1 | -0.071 | 4.528 | 0.034 |
| Matthews<br>Exp.1<br>Periphery | Confidence<br>2 | 252 | 252 | 0.364 | 0.636 | 250 | 2.394 | -1 | -0.009 | 0.938 | 0.334 |
| Matthews<br>Exp.1<br>Fovea | Visibility-Accuracy<br>coupling1 | 252 | 252 | 0.286 | 0.714 | 250 | 1.193 | -1 | -0.001 | 0.234 | 0.629 |
| Matthews<br>Exp.1<br>Periphery | Visibility-Accuracy<br>coupling2 | 252 | 252 | 0.262 | 0.738 | 250 | 1.062 | -1 | 0.000 | -0.001 | 1.000 |
| Matthews<br>Exp.2<br>Fovea | Accuracy1 | 234 | 234 | 0.287 | 0.713 | 232 | 1.785 | -1 | -0.002 | 0.250 | 0.618 |
| Matthews<br>Exp.2<br>Periphery | Accuracy2 | 234 | 234 | 0.837 | 0.163 | 232 | 1.440 | -1 | -0.033 | 5.337 | 0.022 |
| Matthews<br>Exp.2<br>Fovea | Confidence<br>1 | 234 | 234 | 0.351 | 0.649 | 232 | 3.756 | -1 | -0.013 | 0.833 | 0.362 |
| Matthews<br>Exp.2<br>Periphery | Confidence<br>2 | 234 | 234 | 0.992 | 0.008 | 232 | 2.954 | -1 | -0.144 | 11.803 | 0.001 |
| Matthews<br>Exp.2<br>Fovea | Confidence<br>-Accuracy<br>coupling1 | 234 | 234 | 0.264 | 0.736 | 232 | 1.957 | -1 | 0.000 | 0.017 | 0.897 |
| Matthews<br>Exp.2<br>Periphery | Confidence<br>-Accuracy<br>coupling2 | 234 | 234 | 0.923 | 0.077 | 232 | 1.781 | -1 | -0.053 | 7.052 | 0.008 |
| Mei et al.<br>Exp.1 | Accuracy | 560 | 560 | 0.676 | 0.324 | 558 | 2.572 | -1 | -0.016 | 3.488 | 0.062 |
| Mei et al.<br>Exp.1 | Confidence | 560 | 560 | 0.905 | 0.095 | 558 | 2.926 | -1 | -0.034 | 6.530 | 0.011 |
| Mei et al.<br>Exp.1 | Confidence<br>-Accuracy<br>coupling | 560 | 560 | 0.595 | 0.405 | 558 | 1.949 | -1 | -0.010 | 2.789 | 0.095 |
| Mei et al.<br>Exp.1 | Visibility-Accuracy<br>coupling | 560 | 560 | 0.278 | 0.722 | 558 | 1.867 | -1 | 0.000 | 0.117 | 0.733 |
| Mei et al.<br>Exp.1 | Visibility | 560 | 560 | 0.294 | 0.706 | 558 | 4.479 | -1 | -0.002 | 0.275 | 0.600 |
| Mei et al.<br>Exp.2 | Accuracy | 600 | 600 | 0.266 | 0.734 | 598 | 2.220 | -1 | 0.000 | -0.001 | 1.000 |
| Mei et al.<br>Exp.2 | Confidence | 600 | 600 | 0.977 | 0.023 | 598 | 2.497 | -1 | -0.039 | 9.548 | 0.002 |
| Mei et al.<br>Exp.2 | Confidence<br>-Accuracy<br>coupling | 600 | 600 | 0.266 | 0.734 | 598 | 2.143 | -1 | 0.000 | 0.000 | 1.000 |
| Mei et al.<br>Exp.2 | Visibility-Accuracy<br>coupling | 600 | 600 | 0.657 | 0.343 | 598 | 2.740 | -1 | -0.015 | 3.315 | 0.069 |
| Mei et al.<br>Exp.2 | Visibility | 600 | 600 | 0.929 | 0.071 | 598 | 6.627 | -1 | -0.079 | 7.165 | 0.008 |
| Siedlecka<br>Feedback | Accuracy | 3025 | 3025 | 1.000 | 0.000 | 3023 | 24.565 | -1 | -0.160 | 19.754 | 0.000 |
| Siedlecka<br>Feedback | Confidence | 3025 | 3025 | 1.000 | 0.000 | 3023 | 86.358 | -1 | -9.540 | 375.314 | 0.000 |
| Siedlecka<br>Feedback | Confidence<br>-Accuracy<br>coupling | 3025 | 3025 | 1.000 | 0.000 | 3023 | 31.998 | -1 | -1.630 | 162.194 | 0.000 |
| Siedlecka<br>Feedback | Visibility | 3025 | 3025 | 1.000 | 0.000 | 3023 | 92.279 | -1 | -9.524 | 347.781 | 0.000 |
| Del Pin<br>Exp.1 | Accuracy | 270 | 270 | 0.473 | 0.527 | 268 | 2.190 | -1 | -0.015 | 1.832 | 0.177 |
| Del Pin<br>Exp.1 | Visibility-Accuracy<br>coupling | 270 | 270 | 0.263 | 0.737 | 268 | 2.388 | -1 | 0.000 | -0.001 | 1.000 |
| Del Pin<br>Exp.1 | Visibility | 270 | 270 | 0.469 | 0.531 | 268 | 2.452 | -1 | -0.016 | 1.797 | 0.181 |
| Del Pin<br>Exp.2 | Accuracy | 285 | 285 | 0.263 | 0.737 | 283 | 2.056 | -1 | 0.000 | 0.000 | 1.000 |
| Del Pin<br>Exp.2 | Visibility-Accuracy<br>coupling | 285 | 285 | 0.263 | 0.737 | 283 | 1.937 | -1 | 0.000 | 0.000 | 1.000 |
| Del Pin<br>Exp.2 | Visibility | 285 | 285 | 0.705 | 0.295 | 283 | 2.603 | -1 | -0.034 | 3.783 | 0.053 |
| Siedlecka<br>No<br>Feedback | Accuracy | 3400 | 3400 | 0.268 | 0.732 | 3398 | 26.425 | -1 | 0.000 | -0.009 | 1.000 |
| Siedlecka<br>No<br>Feedback | Confidence | 3400 | 3400 | 1.000 | 0.000 | 3398 | 110.424 | -1 | -8.841 | 295.648 | 0.000 |
| Siedlecka<br>No<br>Feedback | Confidence<br>-Accuracy<br>coupling | 3400 | 3400 | 1.000 | 0.000 | 3398 | 36.170 | -1 | -0.337 | 31.910 | 0.000 |
| Siedlecka<br>No<br>Feedback | Visibility | 3400 | 3400 | 1.000 | 0.000 | 3398 | 111.291 | -1 | -9.292 | 309.479 | 0.000 |
| Samaha<br>Perceptual | Accuracy | 1120 | 1120 | 0.697 | 0.303 | 1118 | 7.519 | -1 | -0.025 | 3.679 | 0.055 |
| Samaha<br>Perceptual | Confidence | 1120 | 1120 | 1.000 | 0.000 | 1118 | 20.346 | -1 | -1.377 | 81.104 | 0.000 |
| Samaha<br>Perceptual | Confidence<br>-Accuracy<br>coupling | 1120 | 1120 | 1.000 | 0.000 | 1118 | 9.641 | -1 | -0.352 | 42.318 | 0.000 |
| Samaha<br>WM | Accuracy | 480 | 480 | 0.280 | 0.720 | 478 | 4.144 | -1 | -0.001 | 0.139 | 0.709 |
| Samaha<br>WM | Confidence | 480 | 480 | 1.000 | 0.000 | 478 | 4.950 | -1 | -0.328 | 33.905 | 0.000 |
| Samaha<br>WM | Confidence<br>-Accuracy<br>coupling | 480 | 480 | 1.000 | 0.000 | 478 | 3.990 | -1 | -0.173 | 21.591 | 0.000 |
| Sand &<br>Nilsson | Accuracy | 744 | 744 | 0.267 | 0.733 | 742 | 4.909 | -1 | 0.000 | 0.000 | 1.000 |
| Sand &<br>Nilsson | Visibility-Accuracy<br>coupling | 744 | 744 | 0.267 | 0.733 | 742 | 4.844 | -1 | 0.000 | 0.000 | 1.000 |
| Sand &<br>Nilsson | Visibility | 744 | 744 | 1.000 | 0.000 | 742 | 8.708 | -1 | -0.574 | 52.311 | 0.000 |
| Xue et al.<br>Vertex<br>control | Accuracy | 350 | 350 | 0.354 | 0.646 | 348 | 0.805 | -1 | -0.002 | 0.834 | 0.362 |
| Xue et al.<br>Vertex<br>control | Confidence | 350 | 350 | 1.000 | 0.000 | 348 | 6.267 | -1 | -0.799 | 50.711 | 0.000 |
| Xue et al.<br>Vertex<br>control | Confidence<br>-Accuracy<br>coupling | 350 | 350 | 0.996 | 0.004 | 348 | 1.058 | -1 | -0.039 | 13.456 | 0.000 |
| Xue et al.<br>PFC | Accuracy | 875 | 875 | 0.267 | 0.733 | 873 | 1.850 | -1 | 0.000 | -0.002 | 1.000 |
| Xue et al.<br>PFC | Confidence | 875 | 875 | 1.000 | 0.000 | 873 | 17.877 | -1 | -0.713 | 36.222 | 0.000 |
| Xue et al.<br>PFC | Confidence<br>-Accuracy<br>coupling | 875 | 875 | 1.000 | 0.000 | 873 | 3.288 | -1 | -0.069 | 18.634 | 0.000 |

Note: Two measures of model fits are presented: weighted AICc for linear (w\_lin\_AICc) and exponential (w\_exp\_AICc) models, respectively, and anova test between two models. The larger w\_AICc value represents the supported model. Strong support for one model is provided when w\_AICc > 0.9. Besides, exponential models explain significantly more variances than linear models when anova test yields significant results ( $P < 0.05$ ). Otherwise, insignificant results mean that exponential models cannot outperform linear models in explained variance despite estimating one more parameter.

**Table S2.** Parameter estimates of exponential decay function of visibility data.

| Paper | Experiment | Parameter | $\beta$ | SE | t | p | Sample Size |
| --- | --- | --- | --- | --- | --- | --- | --- |
| Zheng et al. (2024) | Baseline | A | 0.406 | 0.017 | 23.879 | < 0.001 | 59 |
| Zheng et al. (2024) | Baseline | lam | 0.044 | 0.005 | 8.078 | < 0.001 | 59 |
| Zheng et al. (2024) | Baseline | b | 0.057 | 0.014 | 3.984 | < 0.001 | 59 |
| Zheng et al. (2024) | Vis/inv context | A | 0.320 | 0.053 | 6.014 | < 0.001 | 59 |
| Zheng et al. (2024) | Vis/inv context | lam | 0.059 | 0.021 | 2.812 | 0.005 | 59 |
| Zheng et al. (2024) | Vis/inv context | b | 0.140 | 0.023 | 6.078 | < 0.001 | 59 |
| Sand & Nilsson (2017) | Exp.1 | A | 0.228 | 0.028 | 8.105 | < 0.001 | 31 |
| Sand & Nilsson (2017) | Exp.1 | lam | 0.210 | 0.040 | 5.236 | < 0.001 | 31 |
| Sand & Nilsson (2017) | Exp.1 | b | 0.044 | 0.006 | 7.451 | < 0.001 | 31 |
| Jachs et al. (2015) | Exp.1 | A | 0.136 | 0.051 | 2.660 | 0.008 | 10 |
| Jachs et al. (2015) | Exp.1 | lam | 0.426 | 0.193 | 2.209 | 0.028 | 10 |
| Jachs et al. (2015) | Exp.1 | b | 0.029 | 0.004 | 6.649 | < 0.001 | 10 |
| Jachs et al. (2015) | Exp.2 | A | 0.225 | 0.019 | 11.868 | < 0.001 | 13 |
| Jachs et al. (2015) | Exp.2 | lam | 0.087 | 0.020 | 4.295 | < 0.001 | 13 |
| Jachs et al. (2015) | Exp.2 | b | -0.009 | 0.014 | -0.638 | 0.523 | 13 |
| Jachs et al. (2015) | Exp.3 | A | 0.327 | 0.066 | 4.975 | < 0.001 | 10 |
| Jachs et al. (2015) | Exp.3 | lam | 0.366 | 0.093 | 3.936 | < 0.001 | 10 |
| Jachs et al. (2015) | Exp.3 | b | 0.034 | 0.007 | 5.064 | < 0.001 | 10 |
| Mei et al. (2019) | Exp.1 | A | 0.080 | 0.027 | 3.020 | 0.003 | 14 |
| Mei et al. (2019) | Exp.1 | lam | 0.046 | 0.043 | 1.064 | 0.288 | 14 |
| Mei et al. (2019) | Exp.1 | b | 0.026 | 0.033 | 0.789 | 0.430 | 14 |
| Mei et al. (2019) | Exp.2 | A | 0.150 | 0.060 | 2.501 | 0.013 | 15 |
| Mei et al. (2019) | Exp.2 | lam | 0.442 | 0.210 | 2.103 | 0.036 | 15 |
| Mei et al. (2019) | Exp.2 | b | 0.048 | 0.005 | 9.775 | < 0.001 | 15 |
| Siedlecka et al. (2020) | Feedback | A | 0.398 | 0.016 | 25.539 | < 0.001 | 31 |
| Siedlecka et al.<br>(2020) | Feedback | lam | 0.061 | 0.004 | 13.569 | < 0.001 | 31 |
| Siedlecka et al.<br>(2020) | Feedback | b | -0.041 | 0.005 | -8.245 | < 0.001 | 31 |
| Siedlecka et al.<br>(2020) | No feedback | A | 0.353 | 0.017 | 21.394 | < 0.001 | 35 |
| Siedlecka et al.<br>(2020) | No feedback | lam | 0.067 | 0.006 | 12.036 | < 0.001 | 35 |
| Siedlecka et al.<br>(2020) | No feedback | b | -0.007 | 0.005 | -1.465 | 0.143 | 35 |
| Del Pin et al.<br>(2020) | Exp.1 | A | 0.085 | 0.088 | 0.967 | 0.335 | 18 |
| Del Pin et al.<br>(2020) | Exp.1 | lam | 0.085 | 0.096 | 0.884 | 0.378 | 18 |
| Del Pin et al.<br>(2020) | Exp.1 | b | 0.028 | 0.008 | 3.646 | < 0.001 | 18 |
| Del Pin et al.<br>(2020) | Exp.2 | A | 0.017 | 0.177 | 0.097 | 0.923 | 19 |
| Del Pin et al.<br>(2020) | Exp.2 | lam | -0.087 | 0.495 | -0.177 | 0.860 | 19 |
| Del Pin et al.<br>(2020) | Exp.2 | b | -0.018 | 0.208 | -0.085 | 0.932 | 19 |

**Table S3.** Parameter estimates of exponential decay function of confidence data.

| **Paper** | **Experiment** | **Parameter** | **$\beta$** | **SE** | **t** | **p** | **Sample size** |
| --- | --- | --- | --- | --- | --- | --- | --- |
| Jachs et al. (2015) | Exp.1 | A | 0.141 | 0.063 | 2.214 | 0.027 | 10 |
| Jachs et al. (2015) | Exp.1 | lam | 0.557 | 0.276 | 2.021 | 0.044 | 10 |
| Jachs et al. (2015) | Exp.1 | b | 0.021 | 0.004 | 5.458 | < 0.001 | 10 |
| Jachs et al. (2015) | Exp.2 | A | 0.238 | 0.040 | 5.945 | < 0.001 | 13 |
| Jachs et al. (2015) | Exp.2 | lam | 0.378 | 0.080 | 4.753 | < 0.001 | 13 |
| Jachs et al. (2015) | Exp.2 | b | 0.029 | 0.004 | 7.463 | < 0.001 | 13 |
| Jachs et al. (2015) | Exp.3 | A | 0.355 | 0.137 | 2.587 | 0.010 | 10 |
| Jachs et al. (2015) | Exp.3 | lam | 0.848 | 0.297 | 2.851 | 0.005 | 10 |
| Jachs et al. (2015) | Exp.3 | b | 0.014 | 0.004 | 3.166 | 0.002 | 10 |
| Matthews et al. (2018) | Exp.1 fovea | A | 0.158 | 0.035 | 4.465 | < 0.001 | 7 |
| Matthews et al. (2018) | Exp.1 fovea | lam | 0.110 | 0.060 | 1.822 | 0.070 | 7 |
| Matthews et al. (2018) | Exp.1 fovea | b | 0.049 | 0.020 | 2.432 | 0.016 | 7 |
| Matthews et al. (2018) | Exp.1 periphery | A | 0.082 | 0.113 | 0.722 | 0.471 | 7 |
| Matthews et al. (2018) | Exp.1 periphery | lam | 0.580 | 0.872 | 0.665 | 0.506 | 7 |
| Matthews et al. (2018) | Exp.1 periphery | b | 0.080 | 0.007 | 11.592 | < 0.001 | 7 |
| Matthews et al. (2018) | Exp.2 fovea | A | 0.119 | 0.036 | 3.259 | 0.001 | 9 |
| Matthews et al. (2018) | Exp.2 fovea | lam | 0.095 | 0.100 | 0.957 | 0.340 | 9 |
| Matthews et al. (2018) | Exp.2 fovea | b | 0.080 | 0.043 | 1.870 | 0.063 | 9 |
| Matthews et al. (2018) | Exp.2 periphery | A | 0.252 | 0.144 | 1.754 | 0.081 | 9 |
| Matthews et al. (2018) | Exp.2 periphery | lam | 0.690 | 0.403 | 1.712 | 0.088 | 9 |
| Matthews et al. (2018) | Exp.2 periphery | b | 0.080 | 0.008 | 9.673 | < 0.001 | 9 |
| Mei et al. (2019) | Exp.1 | A | 0.326 | 0.710 | 0.459 | 0.646 | 14 |
| Mei et al. (2019) | Exp.1 | lam | 1.798 | 2.102 | 0.855 | 0.393 | 14 |
| Mei et al. (2019) | Exp.1 | b | 0.028 | 0.003 | 8.798 | < 0.001 | 14 |
| Mei et al. (2019) | Exp.2 | A | 0.085 | 0.026 | 3.222 | 0.001 | 15 |
| Mei et al. (2019) | Exp.2 | lam | 0.314 | 0.129 | 2.430 | 0.015 | 15 |
| Mei et al. (2019) | Exp.2 | b | 0.001 | 0.003 | 0.182 | 0.856 | 15 |
| Samaha & Postle<br>(2017) | Perceptual | A | 0.368 | 0.057 | 6.419 | < 0.001 | 16 |
| Samaha & Postle<br>(2017) | Perceptual | lam | 0.352 | 0.067 | 5.230 | < 0.001 | 16 |
| Samaha & Postle<br>(2017) | Perceptual | b | 0.041 | 0.004 | 9.530 | < 0.001 | 16 |
| Samaha & Postle<br>(2017) | WM | A | 0.605 | 0.290 | 2.088 | 0.037 | 16 |
| Samaha & Postle<br>(2017) | WM | lam | 1.227 | 0.428 | 2.869 | 0.004 | 16 |
| Samaha & Postle<br>(2017) | WM | b | 0.016 | 0.005 | 3.382 | 0.001 | 16 |

**Table S4.** Parameter estimates of linear function of accuracy data.

| **Paper** | **Experiment** | **Parameter** | **Estimate** | **SE** | **t** | **p** | **Sam**<br>**ple**<br>**Size** |
| --- | --- | --- | --- | --- | --- | --- | --- |
| Zheng et al. (2024) | Baseline | Intercept | 0.055 | 0.006 | 9.860 | < 0.001 | 59 |
| Zheng et al. (2024) | Baseline | Slope | -0.001 | < 0.001 | -3.832 | < 0.001 | 59 |
| Zheng et al. (2024) | Vis/inv context | Intercept | 0.055 | 0.013 | 4.123 | < 0.001 | 59 |
| Zheng et al. (2024) | Vis/inv context | Slope | < 0.001 | < 0.001 | -1.095 | 0.274 | 59 |
| Sand & Nilsson<br>(2017) | Exp.1 | Intercept | 0.009 | 0.006 | 1.506 | 0.132 | 31 |
| Sand & Nilsson<br>(2017) | Exp.1 | Slope | < 0.001 | < 0.001 | -0.141 | 0.888 | 31 |
| Jachs et al. (2015) | Exp.1 | Intercept | 0.011 | 0.007 | 1.469 | 0.143 | 10 |
| Jachs et al. (2015) | Exp.1 | Slope | < 0.001 | < 0.001 | 0.351 | 0.726 | 10 |
| Jachs et al. (2015) | Exp.2 | Intercept | 0.012 | 0.006 | 2.139 | 0.033 | 13 |
| Jachs et al. (2015) | Exp.2 | Slope | -0.001 | < 0.001 | -2.752 | 0.006 | 13 |
| Jachs et al. (2015) | Exp.3 | Intercept | 0.010 | 0.007 | 1.302 | 0.194 | 10 |
| Jachs et al. (2015) | Exp.3 | Slope | < 0.001 | < 0.001 | -0.399 | 0.690 | 10 |
| Matthews et al.<br>(2018) | Exp.1 fovea | Intercept | 0.022 | 0.010 | 2.185 | 0.030 | 7 |
| Matthews et al.<br>(2018) | Exp.1 fovea | Slope | < 0.001 | < 0.001 | -0.496 | 0.620 | 7 |
| Matthews et al.<br>(2018) | Exp.1 periphery | Intercept | 0.003 | 0.009 | 0.304 | 0.761 | 7 |
| Matthews et al.<br>(2018) | Exp.1 periphery | Slope | < 0.001 | < 0.001 | 0.677 | 0.499 | 7 |
| Matthews et al.<br>(2018) | Exp.2 fovea | Intercept | < 0.001 | 0.012 | 0.026 | 0.979 | 9 |
| Matthews et al.<br>(2018) | Exp.2 fovea | Slope | 0.001 | 0.001 | 1.103 | 0.271 | 9 |
| Matthews et al.<br>(2018) | Exp.2 periphery | Intercept | 0.012 | 0.011 | 1.121 | 0.264 | 9 |
| Matthews et al.<br>(2018) | Exp.2 periphery | Slope | -0.001 | 0.001 | -1.700 | 0.091 | 9 |
| Mei et al. (2019) | Exp.1 | Intercept | 0.028 | 0.006 | 4.759 | < 0.001 | 14 |
| Mei et al. (2019) | Exp.1 | Slope | < 0.001 | < 0.001 | -1.876 | 0.061 | 14 |
| Mei et al. (2019) | Exp.2 | Intercept | 0.024 | 0.005 | 4.650 | < 0.001 | 15 |
| Mei et al. (2019) | Exp.2 | Slope | < 0.001 | < 0.001 | -1.995 | 0.047 | 15 |
| Samaha & Postle<br>(2017) | Perceptual | Intercept | -0.002 | 0.005 | -0.476 | 0.634 | 16 |
| Samaha & Postle<br>(2017) | Perceptual | Slope | < 0.001 | < 0.001 | 1.387 | 0.166 | 16 |
| Samaha & Postle<br>(2017) | WM | Intercept | -0.009 | 0.009 | -1.000 | 0.318 | 16 |
| Samaha & Postle<br>(2017) | WM | Slope | < 0.001 | < 0.001 | 0.772 | 0.440 | 16 |
| Siedlecka et al.<br>(2020) | Feedback | Intercept | 0.017 | 0.003 | 5.249 | < 0.001 | 31 |
| Siedlecka et al.<br>(2020) | Feedback | Slope | < 0.001 | < 0.001 | -3.983 | < 0.001 | 31 |
| Siedlecka et al.<br>(2020) | No feedback | Intercept | 0.013 | 0.003 | 4.229 | < 0.001 | 35 |
| Siedlecka et al.<br>(2020) | No feedback | Slope | < 0.001 | < 0.001 | -5.000 | < 0.001 | 35 |
| Del Pin et al. (2020) | Exp.1 | Intercept | < 0.001 | 0.012 | -0.040 | 0.968 | 18 |
| Del Pin et al. (2020) | Exp.1 | Slope | < 0.001 | < 0.001 | -0.985 | 0.325 | 18 |
| Del Pin et al. (2020) | Exp.2 | Intercept | -0.014 | 0.030 | -0.454 | 0.652 | 19 |
| Del Pin et al. (2020) | Exp.2 | Slope | -0.001 | 0.003 | -0.269 | 0.789 | 19 |

**Table S5.**
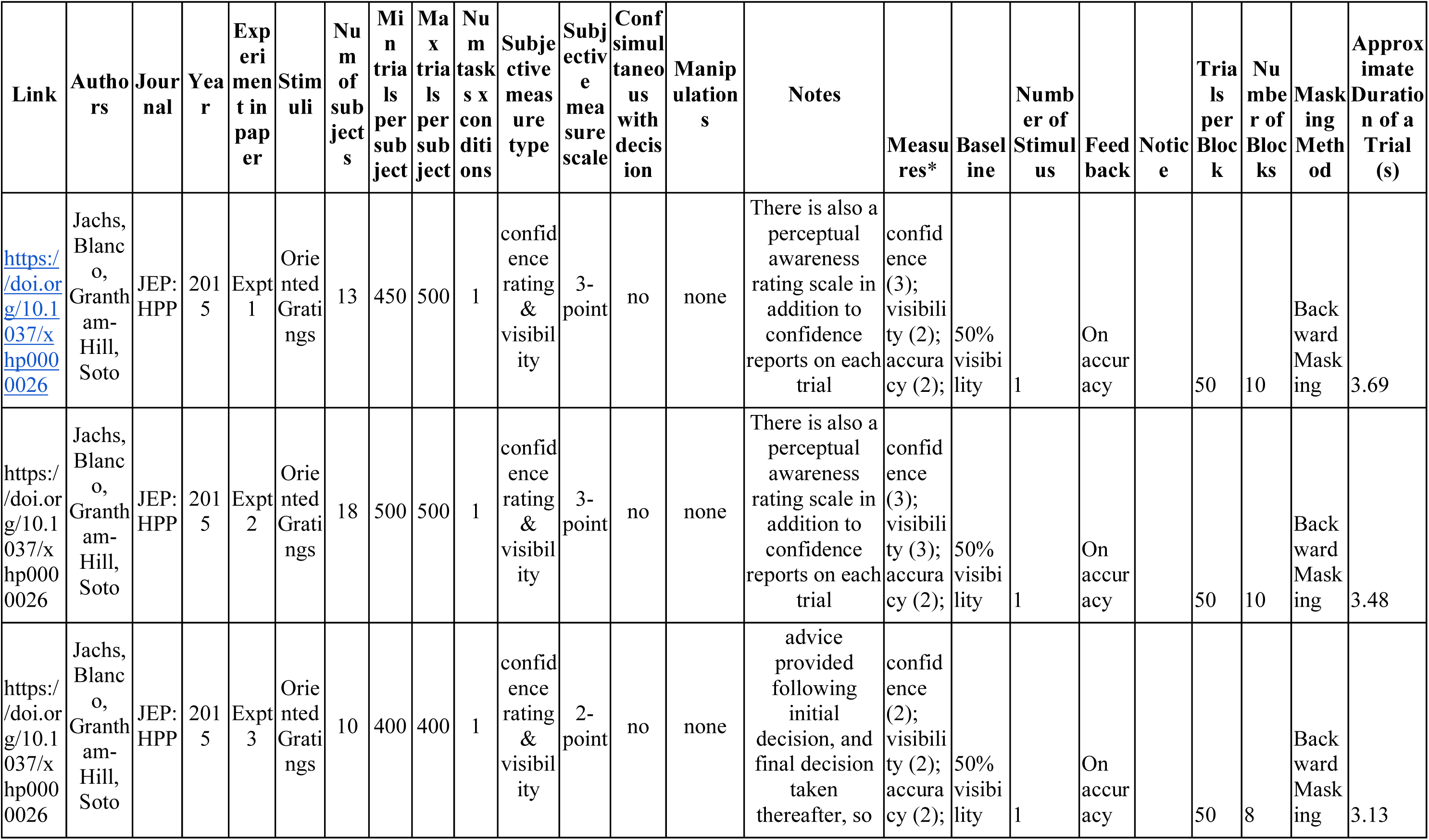
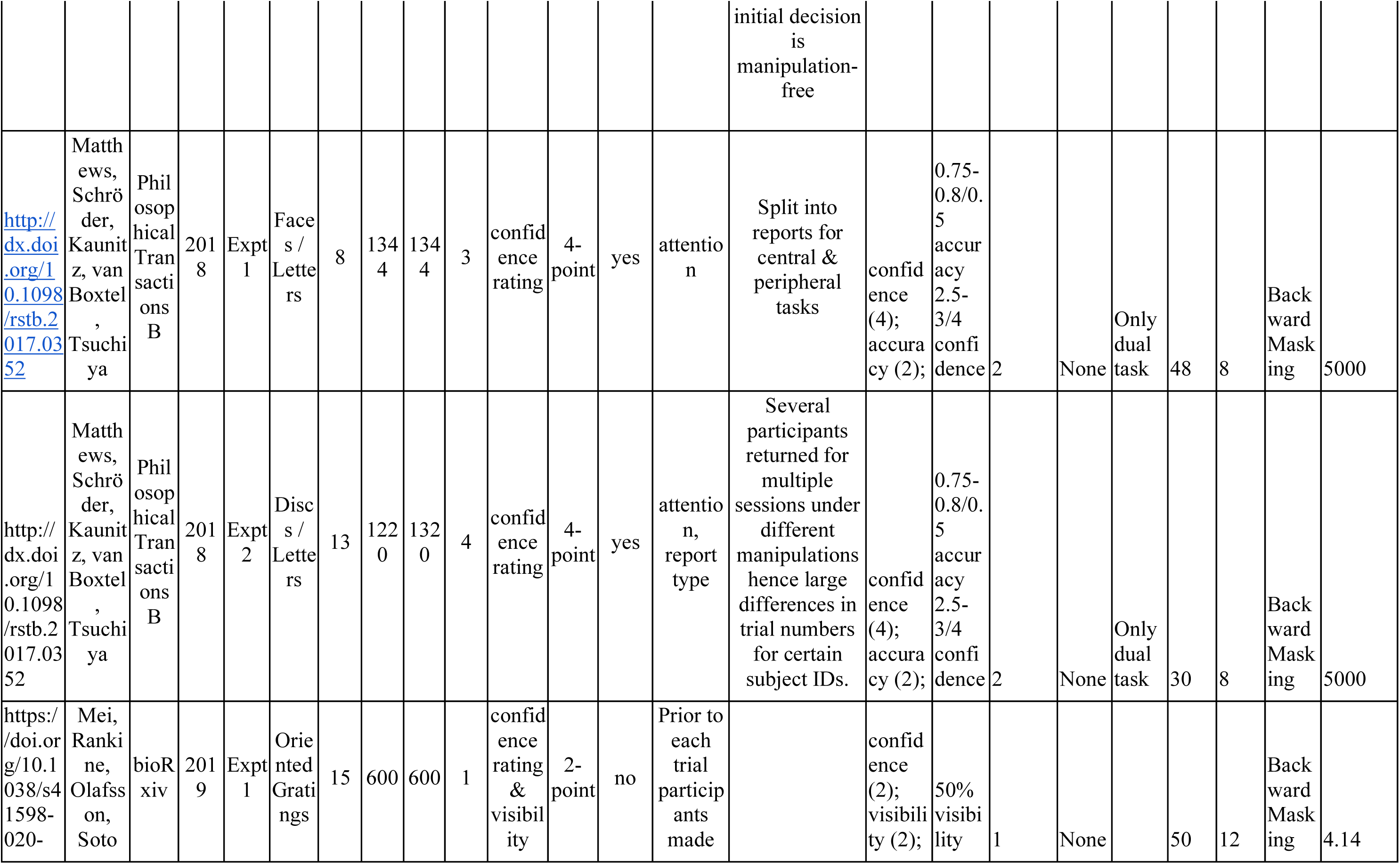
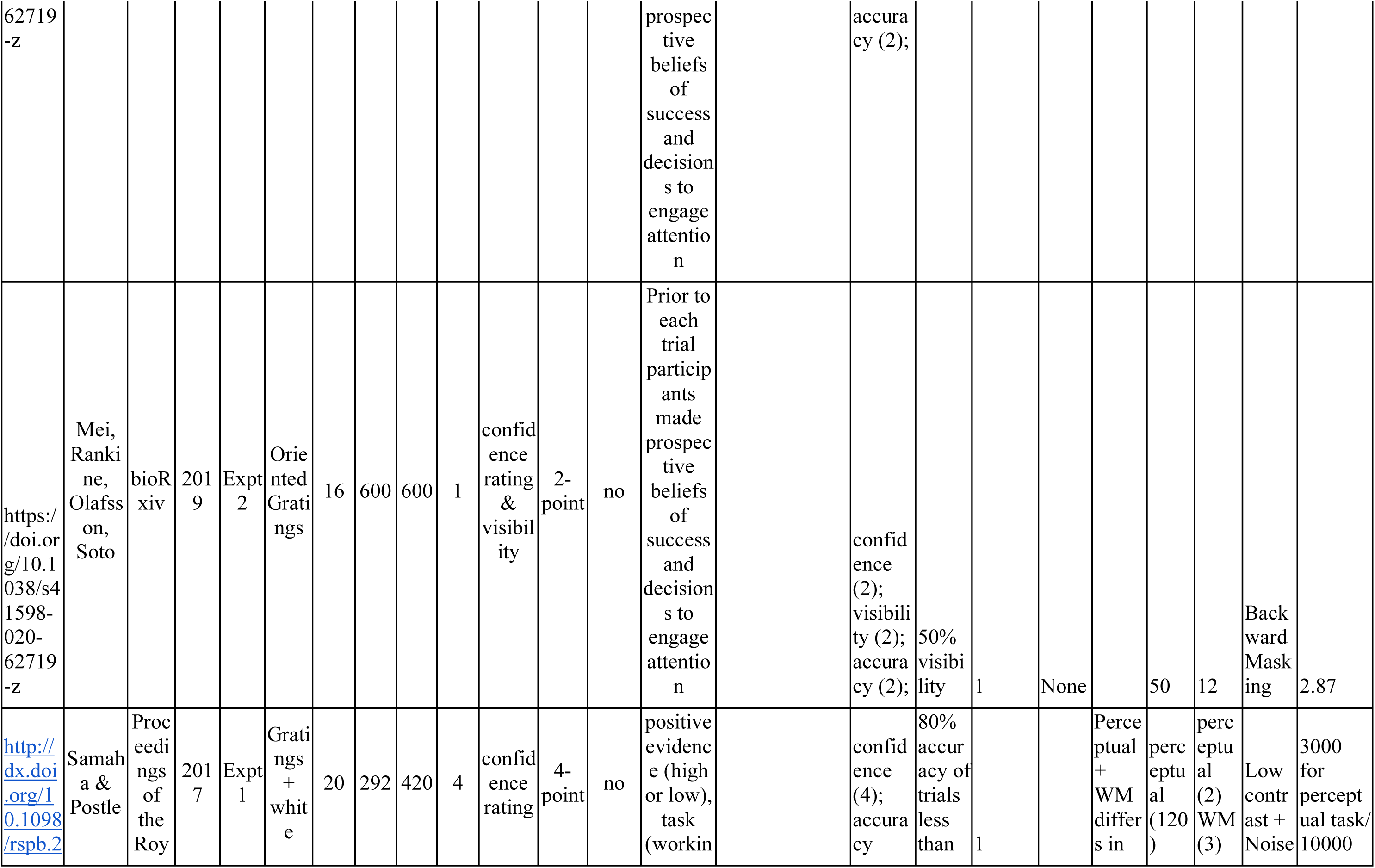
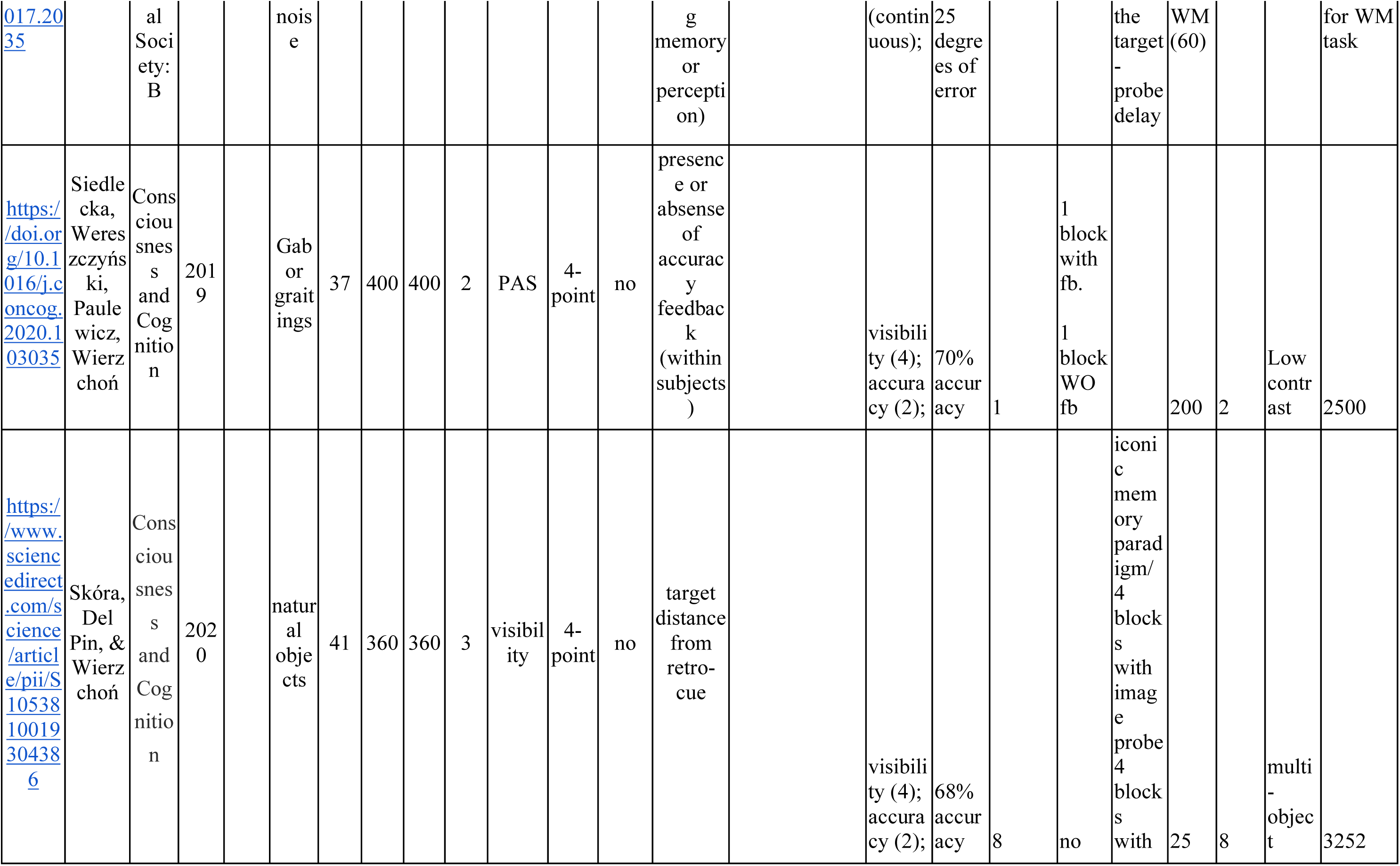
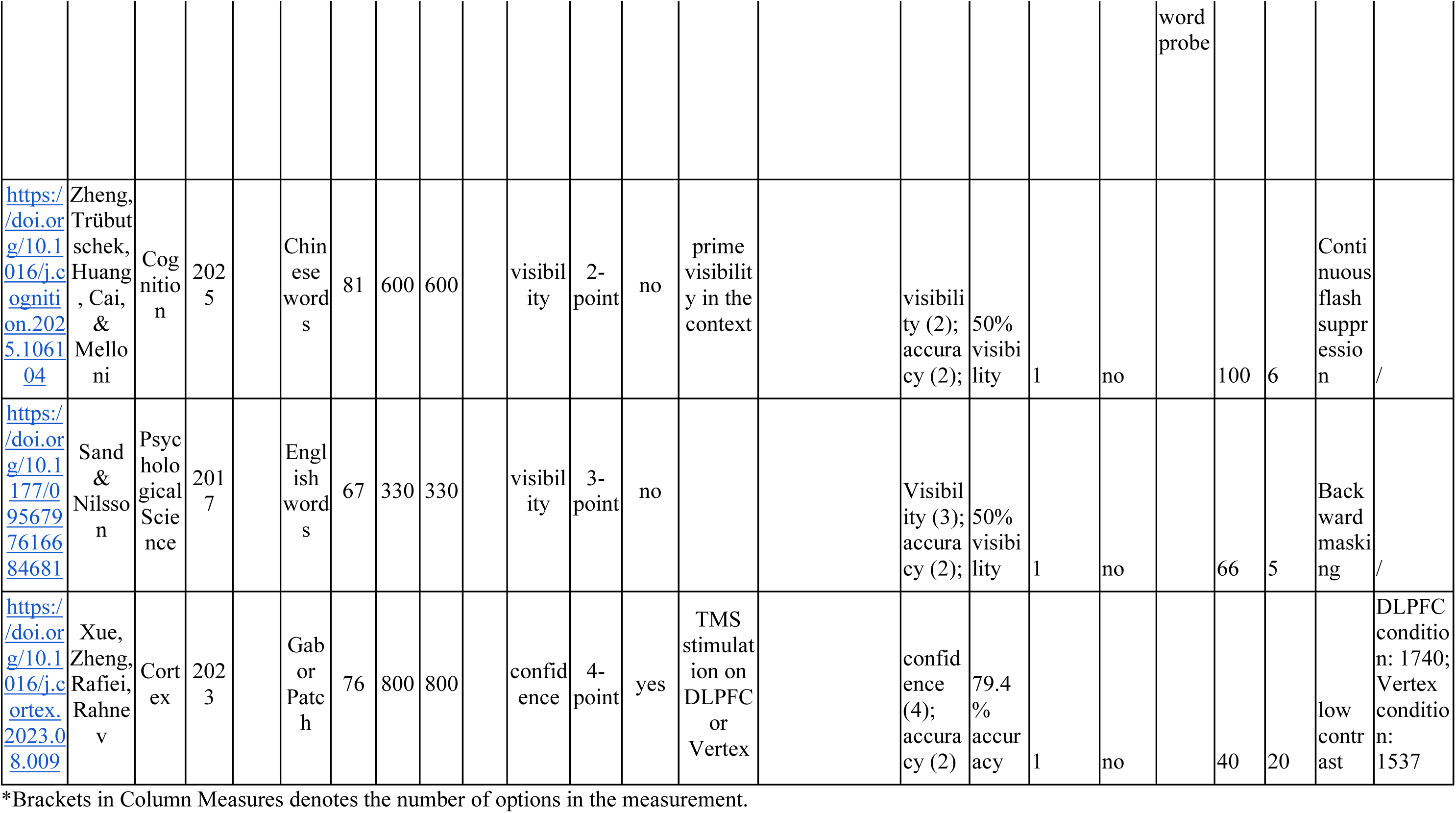
Dataset summary.

**Supplementary Fig. S1.**
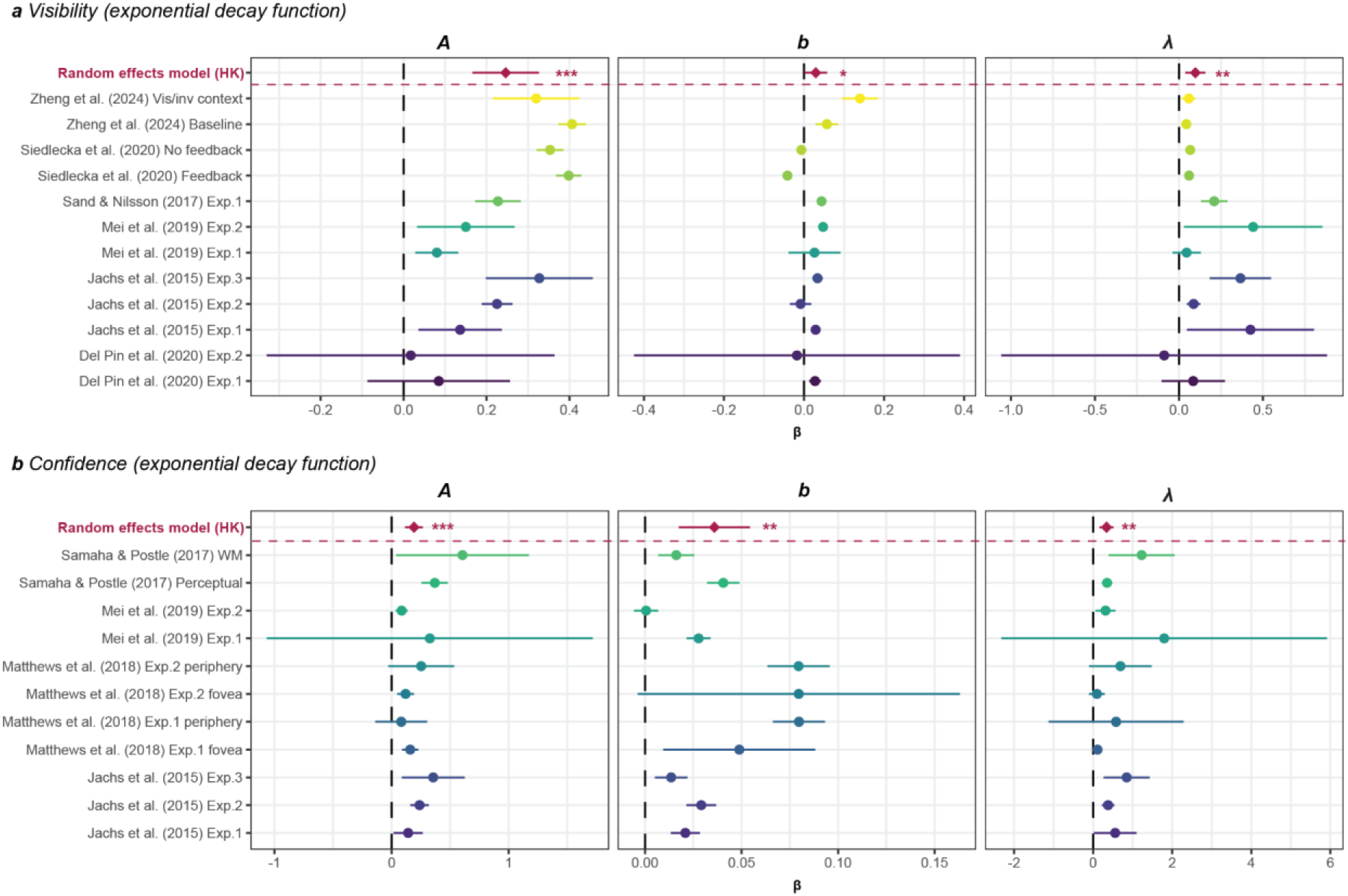
Meta-analytic forest plots of serial dependence parameters for subjective measures. **a**, Visibility, fit with an exponential decay function. **b**, Confidence, fit with the same exponential decay function. Each panel shows the standardized effect size (β) for one fitted parameter of the exponential model: amplitude (**A**), offset (**b**) and decay constant (**λ**). Coloured circles indicate the estimate for each dataset, with horizontal lines showing the corresponding 95% confidence intervals. The maroon diamonds and bars labelled Random effects model (HK) indicate the Hartung–Knapp random-effects meta-analytic estimate and its 95% confidence interval. The vertical dashed black line marks zero effect. Across datasets, both visibility and confidence showed reliably positive amplitudes and decay constants, consistent with robust lag-dependent serial dependence in subjective measures. Visibility also showed a small positive offset, whereas confidence exhibited a clearer positive offset across studies. Asterisks indicate the significance of the pooled effects (* P < 0.05, ** P < 0.01, *** P < 0.001).

**Supplementary Fig. S2.**
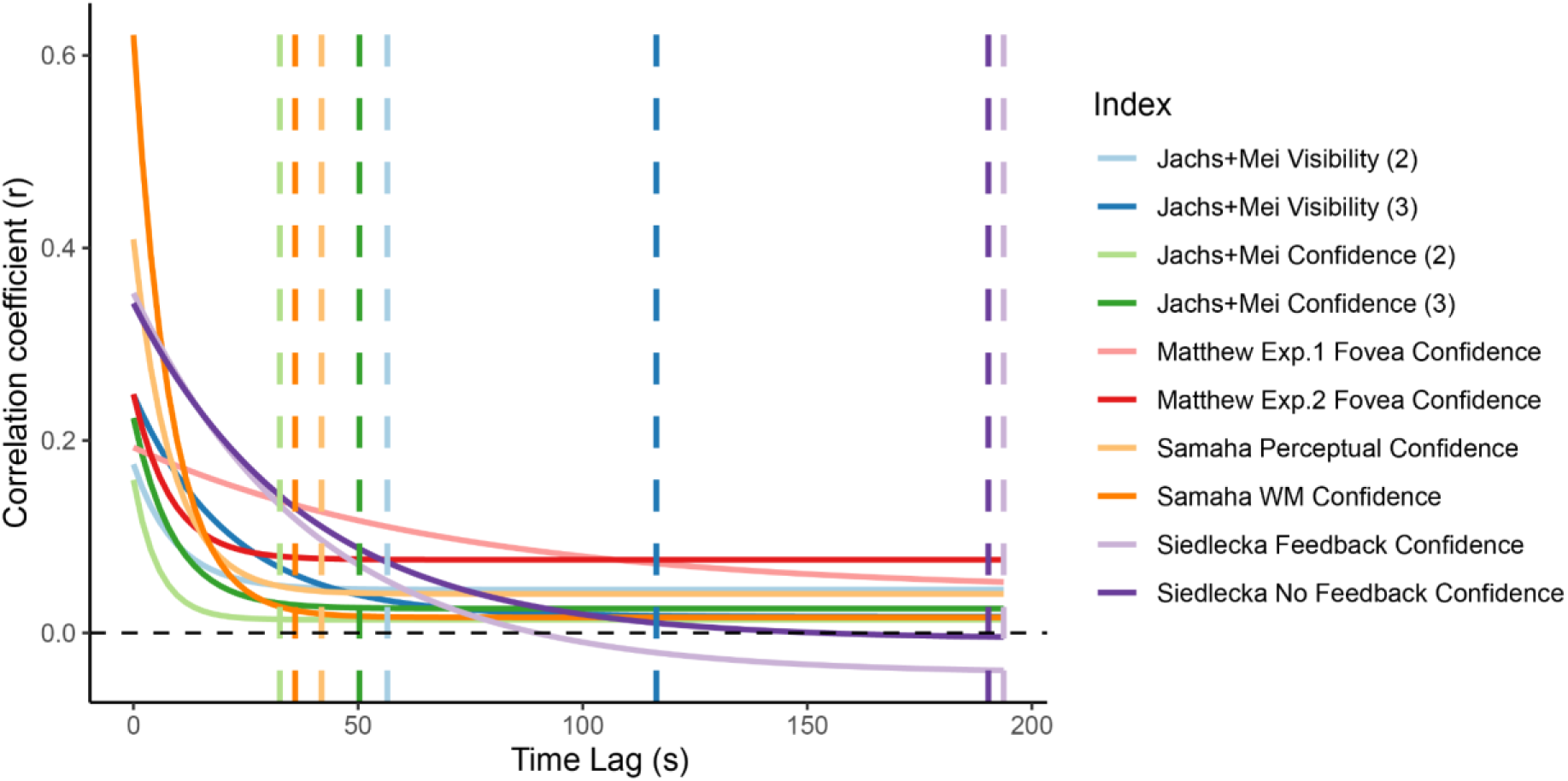
Estimated temporal receptive field of subjective autocorrelation functions in seconds. Exponential fits of autocorrelation functions across datasets show that subjective visibility and confidence exhibit temporal dependencies exceeding 30 s, consistent with prefrontal intrinsic timescales. Vertical bars indicate estimated temporal receptive windows for each dataset, defined as the point at which autocorrelation falls to 1/50 of its initial amplitude.

**Supplementary Fig. S3.**
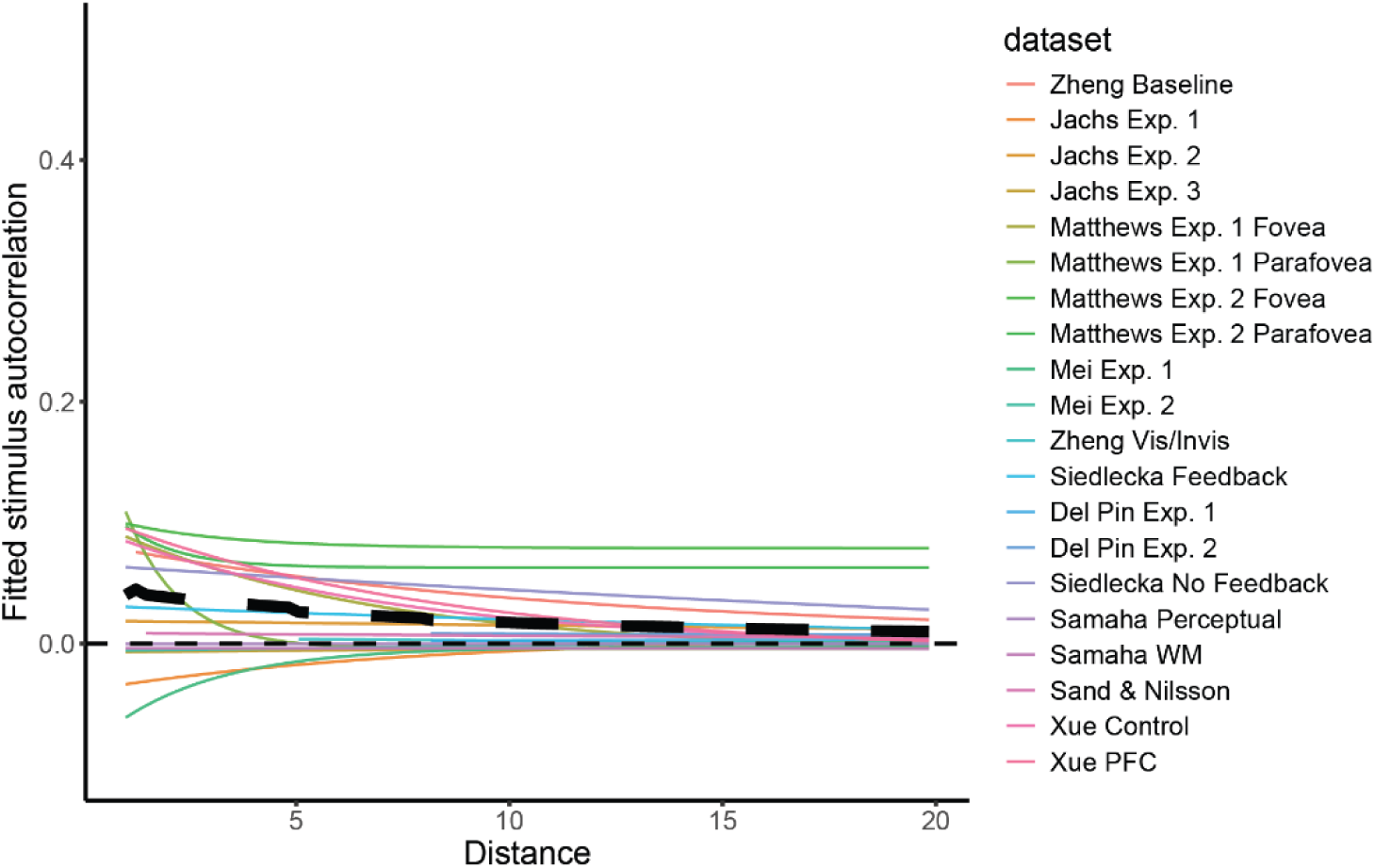
Autocorrelation of first-order stimulus identification responses. Thin coloured lines show fitted autocorrelation functions for the raw identification responses in each dataset; the thick black dashed line shows the weighted average across datasets. Unlike subjective visibility or confidence, the first-order identification responses exhibit only weak autocorrelation overall. This analysis addresses a potential methodological concern: the strong serial dependence observed in subjective measures, and the weaker dependence observed in objective accuracy, could in principle arise because subjective reports are themselves first-order responses, whereas objective accuracy is a second-order quantity derived by matching responses to the true stimulus. To exclude this possibility, we computed the autocorrelation directly on the raw first-order identification responses. The resulting weak effect indicates that the dissociation between subjective and objective measures cannot be reduced to a simple first-order versus second-order difference in how the variables are constructed. Rather, the pronounced serial dependence in subjective measures reflects temporal structure beyond that present in raw identification responses alone.

## References

1. Charles, L., Van Opstal, F., Marti, S. & Dehaene, S. Distinct brain mechanisms for conscious versus subliminal error detection. NeuroImage 73, 80–94 (2013).

2. de Gardelle, V., Charles, L. & Kouider, S. Perceptual awareness and categorical representation of faces: Evidence from masked priming. Consciousness and Cognition 20, 1272–1281 (2011).

3. Dehaene, S. et al. Cerebral mechanisms of word masking and unconscious repetition priming. Nat Neurosci 4, 752–758 (2001).

4. Melloni, L. et al. Synchronization of neural activity across cortical areas correlates with conscious perception. J Neurosci 27, 2858–2865 (2007).

5. Melloni, L., Schwiedrzik, C. M., Müller, N., Rodriguez, E. & Singer, W. Expectations Change the Signatures and Timing of Electrophysiological Correlates of Perceptual Awareness. J. Neurosci. 31, 1386–1396 (2011).

6. Stein, T., Utz, V. & van Opstal, F. Unconscious semantic priming from pictures under backward masking and continuous flash suppression. Consciousness and Cognition 78, 102864 (2020).

7. Stein, T., Kaiser, D., Fahrenfort, J. J. & Gaal, S. van. The human visual system differentially represents subjectively and objectively invisible stimuli. PLOS Biology 19, e3001241 (2021).

8. Zheng, Z., Trübutschek, D., Huang, S., Cai, Y. & Melloni, L. What you saw a while ago determines what you see now: Extending awareness priming to implicit behaviors and uncovering its temporal dynamics. Cognition 259, 106104 (2025).

9. Zheng, Z.-F., Huang, S.-Y., Lu, S. & Cai, Y.-C. Interaction between top-down decision-driven congruency effect and bottom-up input-driven congruency effect is correlated with conscious awareness. Journal of Experimental Psychology: General https://doi.org/10.1037/xge0001483 (2023) doi:10.1037/xge0001483.

10. Koch, C. & Tsuchiya, N. Attention and consciousness: two distinct brain processes. Trends in Cognitive Sciences 11, 16–22 (2007).

11. Lau, H. C. Are We Studying Consciousness Yet? in Frontiers of consciousness (eds Weiskrantz, L. & Davies, M.) 2008–245 (Oxford University Press, 2008).

12. Dellert, T. et al. Dissociating the neural correlates of consciousness and task relevance in face perception using simultaneous EEG-fMRI. J. Neurosci. https://doi.org/10.1523/JNEUROSCI.2799-20.2021 (2021) doi:10.1523/JNEUROSCI.2799-20.2021.

13. Lamme, V. A. F. & Roelfsema, P. R. The distinct modes of vision offered by feedforward and recurrent processing. Trends in Neurosciences 23, 571–579 (2000).

14. Mei, N., Rahnev, D. & Soto, D. Using serial dependence to predict confidence across observers and cognitive domains. Psychon Bull Rev 30, 1596–1608 (2023).

15. Fischer, J. & Whitney, D. Serial dependence in visual perception. Nat Neurosci 17, 738–743 (2014).

16. John-Saaltink, E. S., Kok, P., Lau, H. C. & Lange, F. P. de. Serial Dependence in Perceptual Decisions Is Reflected in Activity Patterns in Primary Visual Cortex. J. Neurosci. 36, 6186–6192 (2016).

17. Kiyonaga, A., Scimeca, J. M., Bliss, D. P. & Whitney, D. Serial Dependence across Perception, Attention, and Memory. Trends in Cognitive Sciences 21, 493–497 (2017).

18. Kandemir, G. & Olivers, C. N. L. Serial dependence is stronger for peripheral than for central vision. Atten Percept Psychophys 88, 44 (2026).

19. Stein, H. et al. Reduced serial dependence suggests deficits in synaptic potentiation in anti-NMDAR encephalitis and schizophrenia. Nat Commun 11, 4250 (2020).

20. Trübutschek, D. & Melloni, L. Stable perceptual phenotype of the magnitude of history biases even in the face of global task complexity. Journal of Vision 23, 1–20 (2023).

21. Snyder, J. S., Schwiedrzik, C. M., Vitela, A. D. & Melloni, L. How previous experience shapes perception in different sensory modalities. Front. Hum. Neurosci. 9, (2015).

22. Verplanck, W. S., Collier, G. H. & Cotton, J. W. Nonindependence of successive responses in measurements of the visual threshold. Journal of Experimental Psychology 44, 273–282 (1952).

23. Koenig, L. & He, B. J. Spontaneous slow cortical potentials and brain oscillations independently influence conscious visual perception. PLOS Biology 23, e3002964 (2025).

24. Monto, S., Palva, S., Voipio, J. & Palva, J. M. Very Slow EEG Fluctuations Predict the Dynamics of Stimulus Detection and Oscillation Amplitudes in Humans. J. Neurosci. 28, 8268–8272 (2008).

25. He, B. J. & Raichle, M. E. The fMRI signal, slow cortical potential and consciousness. Trends in Cognitive Sciences 13, 302–309 (2009).

26. Baria, A. T., Maniscalco, B. & He, B. J. Initial-state-dependent, robust, transient neural dynamics encode conscious visual perception. PLOS Computational Biology 13, e1005806 (2017).

27. Podvalny, E., Flounders, M. W., King, L. E., Holroyd, T. & He, B. J. A dual role of prestimulus spontaneous neural activity in visual object recognition. Nat Commun 10, 3910 (2019).

28. Schwiedrzik, C. M., Singer, W. & Melloni, L. Subjective and objective learning effects dissociate in space and in time. Proceedings of the National Academy of Sciences 108, 4506–4511 (2011).

29. Avneon, M. & Lamy, D. Reexamining unconscious response priming: A liminal-prime paradigm. Consciousness and Cognition 59, 87–103 (2018).

30. Lin, Z. & Murray, S. O. Priming of awareness or how not to measure visual awareness. Journal of Vision 14, 27–27 (2014).

31. Lin, Z. & Murray, S. O. Automaticity of unconscious response inhibition: Comment on Chiu and Aron (2014). Journal of Experimental Psychology: General 144, 244–254 (2015).

32. Rahnev, D., Koizumi, A., McCurdy, L. Y., D’Esposito, M. & Lau, H. Confidence Leak in Perceptual Decision Making. Psychol Sci 26, 1664–1680 (2015).

33. Aguilar-Lleyda, D., Konishi, M., Sackur, J. & de Gardelle, V. Confidence can be automatically integrated across two visual decisions. Journal of Experimental Psychology: Human Perception and Performance 47, 161–171 (2021).

34. Kantner, J., Solinger, L. A., Grybinas, D. & Dobbins, I. G. Confidence carryover during interleaved memory and perception judgments. Mem Cogn 47, 195–211 (2019).

35. Vloeberghs, R., Navarrete Orejudo, L., Urai, A. E. & Desender, K. Trial-by-trial fluctuations in decision criterion shape confidence. Nat Hum Behav https://doi.org/10.1038/s41562-026-02544-y (2026) doi:10.1038/s41562-026-02544-y.

36. Weiskrantz, L. Blindsight revisited. Current Opinion in Neurobiology 6, 215–220 (1996).

37. Balsdon, T. & Azzopardi, P. Absolute and relative blindsight. Consciousness and Cognition 32, 79–91 (2015).

38. Lau, H. C. & Passingham, R. E. Relative blindsight in normal observers and the neural correlate of visual consciousness. PNAS 103, 18763–18768 (2006).

39. Schmid, M. C. et al. Blindsight depends on the lateral geniculate nucleus. Nature 466, 373–377 (2010).

40. Marshall, J. C. & Halligan, P. W. Blindsight and insight in visuo-spatial neglect. Nature 336, 766–767 (1988).

41. Chaudhuri, R., Knoblauch, K., Gariel, M.-A., Kennedy, H. & Wang, X.-J. A Large-Scale Circuit Mechanism for Hierarchical Dynamical Processing in the Primate Cortex. Neuron 88, 419–431 (2015).

42. Murray, J. D. et al. A hierarchy of intrinsic timescales across primate cortex. Nat Neurosci 17, 1661–1663 (2014).

43. Boly, M. et al. Are the Neural Correlates of Consciousness in the Front or in the Back of the Cerebral Cortex? Clinical and Neuroimaging Evidence. J. Neurosci. 37, 9603–9613 (2017).

44. Koch, C., Massimini, M., Boly, M. & Tononi, G. Neural correlates of consciousness: progress and problems. Nat Rev Neurosci 17, 307–321 (2016).

45. Hasson, U. Uncovering a Timescale Hierarchy by Studying the Brain in a Natural Context. J. Neurosci. 45, (2025).

46. Hasson, U., Yang, E., Vallines, I., Heeger, D. J. & Rubin, N. A Hierarchy of Temporal Receptive Windows in Human Cortex. J. Neurosci. 28, 2539–2550 (2008).

47. Honey, C. J. et al. Slow Cortical Dynamics and the Accumulation of Information over Long Timescales. Neuron 76, 423–434 (2012).

48. Gao, R., van den Brink, R. L., Pfeffer, T. & Voytek, B. Neuronal timescales are functionally dynamic and shaped by cortical microarchitecture. eLife 9, e61277 (2020).

49. Samaha, J. & Postle, B. R. Correlated individual differences suggest a common mechanism underlying metacognition in visual perception and visual short-term memory. Proceedings of the Royal Society B: Biological Sciences 284, 20172035 (2017).

50. Siedlecka, M., Wereszczyński, M., Paulewicz, B. & Wierzchoń, M. Visual awareness judgments are sensitive to accuracy feedback in stimulus discrimination tasks. Consciousness and Cognition 86, 103035 (2020).

51. Sand, A. & Nilsson, M. E. When Perception Trumps Reality: Perceived, Not Objective, Meaning of Primes Drives Stroop Priming. Psychol Sci 28, 346–355 (2017).

52. Jachs, B., Blanco, M. J., Grantham-Hill, S. & Soto, D. On the independence of visual awareness and metacognition: A signal detection theoretic analysis. Journal of Experimental Psychology: Human Perception and Performance 41, 269–276 (2015).

53. Mei, N., Rankine, S., Olafsson, E. & Soto, D. Similar history biases for distinct prospective decisions of self-performance. Sci Rep 10, 5854 (2020).

54. Matthews, J., Schröder, P., Kaunitz, L., van Boxtel, J. J. A. & Tsuchiya, N. Conscious access in the near absence of attention: critical extensions on the dual-task paradigm. Philosophical Transactions of the Royal Society B: Biological Sciences 373, 20170352 (2018).

55. Del Pin, S. H., Skóra, Z., Sandberg, K., Overgaard, M. & Wierzchoń, M. Comparing theories of consciousness: Object position, not probe modality, reliably influences experience and accuracy in object recognition tasks. Consciousness and Cognition 84, 102990 (2020).

56. Rahnev, D. et al. The Confidence Database. Nat Hum Behav 4, 317–325 (2020).

57. He, B. J. Scale-free brain activity: past, present, and future. Trends in Cognitive Sciences 18, 480–487 (2014).

58. Ramsøy, T. Z. & Overgaard, M. Perceptual Awareness Scale. 10.1037/t59063-000 (2018).

59. Wierzchoń, M., Anzulewicz, A., Hobot, J., Paulewicz, B. & Sackur, J. In search of the optimal measure of awareness: Discrete or continuous? Consciousness and Cognition 75, 102798 (2019).

60. Fahrenfort, J. J., Johnson, P. A., Kloosterman, N. A., Stein, T. & Gaal, S. van. Criterion placement threatens the construct validity of neural measures of consciousness. 2024.02.22.581517 Preprint at 10.1101/2024.02.22.581517 (2024).

61. Jannati, A. & Di Lollo, V. Relative blindsight arises from a criterion confound in metacontrast masking: Implications for theories of consciousness. Consciousness and Cognition 21, 307–314 (2012).

62. Vloeberghs, R., Urai, A. E., Desender, K. & Linderman, S. W. A Bayesian hierarchical model of trial-to-trial fluctuations in decision criterion. PLOS Computational Biology 21, e1013291 (2025).

63. Bang, J. W. & Rahnev, D. Stimulus expectation alters decision criterion but not sensory signal in perceptual decision making. Scientific Reports 7, 17072 (2017).

64. Xue, K., Zheng, Y., Rafiei, F. & Rahnev, D. The timing of confidence computations in human prefrontal cortex. Cortex 168, 167–175 (2023).

65. Dehaene, S., Changeux, J.-P., Naccache, L., Sackur, J. & Sergent, C. Conscious, preconscious, and subliminal processing: a testable taxonomy. Trends in Cognitive Sciences 10, 204–211 (2006).

66. Dehaene, S. & Changeux, J.-P. Ongoing Spontaneous Activity Controls Access to Consciousness: A Neuronal Model for Inattentional Blindness. PLoS Biol 3, e141 (2005).

67. Raffone, A., Srinivasan, N. & van Leeuwen, C. The interplay of attention and consciousness in visual search, attentional blink and working memory consolidation. Phil. Trans. R. Soc. B 369, 20130215 (2014).

68. Dehaene, S., Kerszberg, M. & Changeux, J.-P. A Neuronal Model of a Global Workspace in Effortful Cognitive Tasks. Proceedings of the National Academy of Sciences of the United States of America 95, 14529–14534 (1998).

69. Mashour, G. A., Roelfsema, P., Changeux, J.-P. & Dehaene, S. Conscious Processing and the Global Neuronal Workspace Hypothesis. Neuron 105, 776–798 (2020).

70. Crick, F. & Koch, C. Towards a neurobiological theory of consciousness. in Seminars in the Neurosciences vol. 2 203 (London, 1990).

71. Mathewson, K. E., Gratton, G., Fabiani, M., Beck, D. M. & Ro, T. To See or Not to See: Prestimulus α Phase Predicts Visual Awareness. J. Neurosci. 29, 2725–2732 (2009).

72. GNU Scientific Library Reference Manual: For GSL Version 1.12. (Network Theory, Bristol, 2009).

73. Balduzzi, S., Rücker, G. & Schwarzer, G. How to perform a meta-analysis with R: a practical tutorial. Evid Based Mental Health 22, 153–160 (2019).

74. Viechtbauer, W. Bias and Efficiency of Meta-Analytic Variance Estimators in the Random-Effects Model. Journal of Educational and Behavioral Statistics 30, 261–293 (2005).

75. Hartung, J. & Knapp, G. On tests of the overall treatment effect in meta-analysis with normally distributed responses. Statistics in Medicine 20, 1771–1782 (2001).

76. Hartung, J. & Knapp, G. A refined method for the meta-analysis of controlled clinical trials with binary outcome. Statistics in Medicine 20, 3875–3889 (2001).

